# An effective cold storage method for stockpiling *Halyomorpha halys* eggs for field surveys and laboratory rearing of *Trissolcus japonicus*

**DOI:** 10.1101/2020.04.30.071183

**Authors:** Warren H. L. Wong, Matt A. Walz, Angela B. Oscienny, Jade L. Sherwood, Paul K. Abram

## Abstract

An effective stockpiling method for egg masses of the invasive brown marmorated stink bug (*Halyomorpha halys* [Stål]; Hemiptera: Pentatomidae) would be useful for rearing and field studies of its egg parasitoid *Trissolcus japonicus* (Ashmead) (Hymenoptera: Scelionidae). The current method of stockpiling *H. halys* egg masses at sub-zero temperatures has lethal and sublethal fitness consequences for *T. japonicus*. We show that parasitoid emergence from egg masses refrigerated at 8°C for up to two months before parasitism is higher than from frozen egg masses and usually has minimal or no sublethal fitness effects (sex ratio, development time, activity, fecundity, longevity, and weight) on emerging *T. japonicus*. Only after two months of host egg refrigeration did the emergence of *T. japonicus* begin to decrease significantly (by 9.6% relative to untreated viable egg masses), whereas egg masses previously frozen at -80°C had a 58.8% reduction in parasitoid emergence after 14 days of storage. Refrigerated egg masses that were subsequently exposed to average field temperatures (warm: 22.9°C; cool: 13.2°C) were still suitable for *T. japonicus* parasitism after 7 days, while viable egg masses exposed to warm temperatures for 7 days before parasitism had 24.1% lower parasitoid emergence. Our results demonstrate that refrigeration at 8°C, while resulting in complete mortality of *H. halys* embryos after 10 days, are more suitable for *T. japonicus* parasitism than those stored at sub-zero temperatures. The quantity and quality of *H. halys* eggs that can be stockpiled with this method could facilitate *T. japonicus* laboratory colony maintenance, field monitoring, and releases.

## Introduction

Cold storage is an important method for extending the shelf life of parasitoid biological control agents and their hosts (Rathee and Ram 2018). On-demand production of parasitoids can be facilitated by stockpiling host material in cool or freezing temperatures, which may either extend or terminate host development (Mahmoud and Lim 2007; Ayvaz and Karabörklü 2008) depending on the temperature and the duration of storage. Different host insect life stages (Ayvaz and Karabörklü 2008; Tang et al. 2014; Forouzan et al. 2018) can be cold-stored across a range of temperatures, including above-freezing (hereafter, “refrigeration”; typically 2 - 10°C) and sub-zero (hereafter “freezing”; as low as -80°C) temperatures (Mainali and Lim 2013; Spínola-Filho et al. 2014; McIntosh et al. 2019). Cold storage durations for host material in past studies have varied from relatively short-term (less than 7 days), intermediate amounts of time (weeks to months) or much longer time frames (e.g. > 1 year) (Kivan and Kilic 2005; Spínola-Filho et al. 2014; McIntosh et al. 2019). In many cases, cold storage treatments are an effective strategy for extending the period of availability of host material for research or production of parasitoids for releases against pest insects (Orr 1988; Goldson et al. 1993; Rathee and Ram 2018). However, it is important to assess the limitations of using cold-stored host material, as it may negatively impact the quantity or quality of parasitoids that are produced (Orr 1988; Chen and Leopold 2007; Colinet and Boivin 2011).

The consequences of reduced host suitability for parasitoids reared on cold-stored hosts can take the form of (i) reduced acceptance of cold-stored hosts by parasitoids, (ii) lethal effects on developing parasitoids in accepted host eggs, or (iii) sublethal effects on developing parasitoids in accepted host eggs. Lethal effects, whereby cold storage reduces parasitoid offspring emergence, are often due to reduced host quality caused by cold storage (Mahmoud and Lim 2007; Mainali and Lim 2013; McIntosh et al. 2019). Sub-lethal effects of cold storage do not kill immature parasitoids, but subsequently cause changes in their physiology, morphology or behaviour (Mahmoud and Lim 2007; Mainali and Lim 2013; McIntosh et al. 2019). Parasitoid traits that can be affected by cold storage of host material include development time, behaviours related to host acceptance (e.g. oviposition, marking), sex ratio, fecundity, and progeny body size (Kivan and Kilic. 2005; Mahmoud and Lim 2007; Molaei et al. 2015). However, in some cases, cold storage can kill hosts while only having minimal sub-lethal and lethal effects on parasitoids (e.g., Peverieri et al. 2014). This can often be the case for idiobiont egg parasitoids, who by definition develop on dead hosts and are therefore less likely to be affected by reductions in host viability due to cold storage (e.g., Alim and Lim 2009; Peverieri et al. 2014).

The brown marmorated stink bug, *Halyomorpha halys* (Stål) (Hemiptera: Pentatomidae) is an invasive, polyphagous pest that has recently been responsible for economic losses in tree fruit, ornamentals, vegetables and legume crops in North America and Europe (Haye and Weber 2017; Leskey and Nielsen 2018). *Trissolcus japonicus* (Ashmead) (Hymenoptera: Platygastridae) is the egg parasitoid most commonly associated with *H. halys* in Asia (Zhang et al. 2017). Much research effort has been devoted to surveying for native natural enemies of *H. halys* eggs in invaded ranges, as well as detecting the abundance and spread of adventive populations of *T. japonicus* (Haye et al. 2015; Talamas et al. 2015; Herlihy et al. 2016; Abram et al. 2017; Dieckhoff et al. 2017; Abram et al. 2019; Stahl et al. 2019a). Several laboratory studies also continue to be conducted on the host range, behaviour, and basic biology of *T. japonicus* (e.g., Hedstrom et al. 2017; Nystrom Santacruz et al. 2017; Bertoldi et al. 2019; Haye et al. 2019), and inoculative or augmentative biological control releases in several areas are planned, ongoing, or are being considered in several areas (e.g., Lowenstein et al. 2020). A stockpiling technique for *H. halys* eggs for parasitism by *T. japonicus* would be useful for “sentinel egg mass” field studies investigating *T. japonicus* distribution and abundance, as well as laboratory colony maintenance and production of larger numbers of parasitoids for field releases. Currently, deep-freezing *H. halys* egg masses at -80°C is the main method used for stockpiling in support of laboratory and field studies (Abram et al. 2017; McIntosh et al. 2019). In some earlier studies, deep-freezing of *H. halys* eggs was observed to increase the development success of some native European and North American parasitoids, although effects were inconsistent across parasitoid species (Haye et al. 2015; Herlihy et al. 2016; Abram et al. 2017). For *T. japonicus* however, a recent study demonstrated that deep-freezing *H. halys* eggs at -80°C had both lethal and sublethal negative effects on parasitism and parasitoid offspring (McIntosh et al. 2019). Freezing of host eggs reduced emergence by 53% overall and by 1-3% for each additional month in storage (McIntosh et al. 2019). In addition, developing in frozen *H. halys* eggs reduced emergence and delayed development time by 5-6 days compared to fresh eggs (McIntosh et al. 2019). After frozen eggs were removed from the freezer, parasitoid emergence decreased by 5-8% for each day the egg mass subsequently spent at moderate temperatures (mimicking field exposure) before being parasitized by *T. japonicus* (McIntosh et al. 2019). To date, there are no other studies testing lethal and sublethal effects of other, potentially less detrimental cold storage temperatures (i.e., refrigeration) for *H. halys* egg masses for parasitism by *T. japonicus*, despite cold storage being a commonly tested method for other scelionid parasitoids in biological control programs targeting other stink bug pests (Orr 1988; Cingolani et al. 2018).

In this study, we evaluated the effects of refrigerating *H. halys* egg masses at 8°C on subsequent parasitism success by *T. japonicus* and sublethal effects on their offspring. As a starting point, we chose 8°C, which is within a range of temperatures (4-10°C) often used in entomology laboratories to temporarily delay insect development (Paul K. Abram, personal observations), and also because it lies between freezing (0°C) and the lower developmental threshold of *H. halys* (∼12-13°C) (Nielsen et al. 2008; Haye et al. 2014). Our goals were to (1) determine the effect of different refrigeration durations on *H. halys* egg viability (nymph emergence); (2) determine the effect of different refrigeration durations on *T. japonicus* parasitism success and offspring fitness parameters; (3) directly compare refrigeration of *H. halys* eggs at 8°C to the previously used method of freezing *H. halys* eggs at -80°C; (4) evaluate the quality of refrigerated *H. halys* eggs for parasitism by *T. japonicus* at different times after removal from refrigeration; (5) determine the maximum length of time that *H. halys* eggs can be refrigerated at 8°C without a decline in subsequent parasitism levels by *T. japonicus*. Our results demonstrate that refrigerating *H. halys* eggs at 8°C could be useful as a method for stockpiling host material for *T. japonicus* rearing, with applications for conducting research and improving the production of this biological control agent for inoculative and augmentative biological control releases.

## Materials and Methods

### Insect colonies

Adult *H. halys* were collected from the Fraser Valley region of British Columbia, Canada. *Halyomorpha halys* were reared in mesh cages (45 cm^3^) containing potted, greenhouse-grown, bell peppers (*Capsicum annuum* L. [Solanaceae]), cabbage (*Brassica oleracea* L. [Brassicaceae]), and pea (*Pisum sativum* L. [Fabaceae]) plants. Cages were kept in rearing rooms (24°C, 50% RH, 16:8 h L:D) at Agriculture and Agri-Food Canada’s Agassiz Research and Development Centre (ARDC) (Agassiz, British Columbia, Canada). *Halyomorpha halys* were fed a diet of carrots (*Daucus carota sativus*), green beans (*Phaseolus vulgaris*), bananas (*Musa acuminate*), and sunflower seeds (*Helianthus annuus*), with food replaced every three to four days. Potted pepper, cabbage and pea plants were replaced, and the cages cleaned every seven days. Eggs were collected daily from each cage to be used for experiments and parasitoid rearing. Eggs collected once per week were placed into 1 L cylindrical plastic containers with green beans and allowed to hatch and age to approximately 2^nd^ instar. At 2^nd^ instar, nymphs were transferred to the mesh rearing cages containing two potted bell pepper plants and two potted pea plants, as well as carrots and green beans for food. The rearing cage was placed in a climate-controlled growth chamber set to 23°C, 65% RH and 16:8 h L:D until the nymphs were approximately 4^th^ instar, at which point the rearing cage was transferred to a rearing room containing *H. halys* adults.

The colony of *T. japonicus* used in this study was originally collected by the USDA-ARS (Dr. Kim Hoelmer and colleagues) in Beijing, China in 2009. This is the same colony used in several past laboratory studies of *T. japonicus* in North America (Hedstrom et al. 2017; Botch and Delfosse 2018; Kim Holemer, personal communication) and being considered for intentional releases in Canada, the USA, and Europe (PKA, personal observations). Individuals were acquired in 2017 to start a colony in a certified containment facility at the ARDC. To obtain *T. japonicus* for experiments, mated females were isolated 5-14 days after emergence in 1.7 mL microcentrifuge tubes for at least 24 h and provided with a drop of liquid honey on the underside of the lid. Four to five freshly collected *H. halys* egg masses were glued to a strip of paper and placed in a ventilated plastic container (length: 70 mm, width: 70 mm, height: 100 mm) with honey spread over a portion of the mesh lid. Five to six of the isolated *T. japonicus* females were then transferred to these containers for 24 h to parasitize the *H. halys* egg masses, after which the parasitoids were removed from the containers. After completing development, emerging *T. japonicus* adults remained in the containers and were fed a diet of liquid honey spread on the mesh lid of the container and misted with water twice a week. Females used to parasitize *H. halys* eggs masses in all experiments were given adequate time to develop full egg loads (6-10 days, see below) and had been isolated in a 1.7 mL microcentrifuge tube for at least 24 hours before being offered *H. halys* egg masses. All experiments involving live *T. japonicus* were conducted in a containment facility under standard rearing conditions: 22-24°C; humidity 30-50% RH; photoperiod: 16:8 h L:D.

### Experiment 1: Effect of refrigeration on H. halys nymphs

To test the effects of refrigeration on *H. halys* egg survival, egg masses (< 24 h since being laid) were placed in an 8°C incubation chamber for different lengths of time. First, egg masses were divided in half with forceps. After egg masses were divided, each half was glued to a quarter section of 389 grade 45 mm diameter filter paper (Sartorius; Wood Dale, IL, USA), and placed into a 50 × 9 mm Petri dish (Falcon; Durham, NC, USA). Four drops of water were placed on the underside of the lid (∼0.5 µL per drop) to maintain adequate humidity; a preliminary experiment suggested that this maximized *H. halys* emergence levels. Half of each egg mass (13 ± 1 eggs) was placed at room temperature (23 ± 2°C) as a paired control for baseline hatching success of *H. halys* nymphs, while the other half of the same egg mass was placed in a randomly assigned refrigeration treatment (0, 2, 4, 6, 10, or 14 days; n = 20 per treatment). Refrigerated egg masses (following their refrigeration treatment) and paired controls were placed at room temperature for 14 days. After 14 days, when all emergence had concluded, the numbers of emerged nymphs, aborted eggs, and unemerged nymphs were counted. Proportion emergence was calculated as the number of emerged nymphs from the total number of eggs in each egg mass.

### Experiment 2: Effect of refrigeration duration on T. japonicus survival and fitness

The goal of the second experiment was to test the effects of different short-term durations of 8°C refrigeration on the suitability of *H. halys* eggs for *T. japonicus* in terms of both lethal and sublethal effects on parasitoids. Newly collected (< 24 h since being laid) *H. halys* eggs masses were collected and divided in half with forceps. Each half was glued to a quarter section of 389 grade 45 mm diameter filter paper (Satorius; Wood Dale, IL, USA), and placed into a 50 × 9 mm Petri dish (Falcon; Durhan, NC, USA) with a tight-fitting lid. Four drops of water were placed on the underside of the lid (∼0.5 µL per drop), to maintain adequate humidity. One half of each *H. halys* egg mass (mean of 14 ± 1 eggs) was immediately placed at 23 ± 1°C and parasitized by a single *T. japonicus* female for 24 h, while the other half of the same egg mass was placed in a randomly assigned refrigeration duration treatment (0, 2, 4, 6, 10, or 14 days; n = 20 per duration), removed from storage following this incubation period, and then immediately exposed to a different experimental *T. japonicus* female for 24 h at 23 ± 1°C. After being exposed to parasitoids, refrigerated egg masses and paired controls were kept at room temperature for 14 days. The number, development time (calculated as the midpoint between the last two observations), and sex of the emerging *T. japonicus* were recorded daily. Development times were averaged for each egg mass (separately for each sex), and these averages were used in subsequent analysis. Sex ratio was calculated as the proportion of male offspring emerging from each egg mass. Egg masses were dissected to count the number of unemerged adult parasitoids; however, these were not included in calculations of proportion emergence. As we did not directly observe parasitoid behaviour, and some oviposition may have resulted in no detectable offspring development (i.e., if they died before reaching the larval stage), we could not directly assess to what degree differences in parasitoid emergence were due to different levels of acceptance (oviposition) *versus* different levels of immature parasitoid developmental success. However, as parasitism levels in both treatments were very high on average (> 90%; see *Results*), acceptance levels were likely high.

For a subset of emerging parasitoids, we measured four fitness-related behavioural and life history parameters (Roitberg et al. 2001) to assess potential sublethal effects of the different refrigeration durations: locomotor activity, fecundity, longevity, and weight. Due to the large number of specimens involved, and the apparent consistency in quality of the unrefrigerated paired controls (see *Results*), we only measured these parameters from parasitoids emerged from the halves of egg masses that were refrigerated (2-14 days) and those that were in the 0-day refrigeration treatment; that is, we did not include parasitoids from paired controls for refrigerated egg masses.

We used locomotor activity of *T. japonicus* as a behavioural proxy for the intensity of host searching and overall vigor (e.g., Rasmussen et al. 2018). To measure the locomotor activity of female parasitoids emerging from eggs that had been refrigerated for different durations, the first emerged female from each replicate was transferred to a glass tube (diameter: 2.5 cm; length: 12.5 cm) capped at both ends with cellulose plugs (Narrow flugs, Diamed, Canada) with access to two drops of honey (*ad libitum*). Taking into account the partially inserted cellulose plugs, there was ∼34 cm^3^ of space available for parasitoids to move around inside the tube. These tubes were randomly assigned to cells in one of two insect locomotor activity monitors (LAM25H, Trikinetics, Waltham, MA, USA), each with 32 cells arranged in a 4 by 8 grid. These activity monitors recorded the total number of times each insect crossed an infrared beam during each 5 min time interval (288 intervals/day) (Pfeiffenberger et al. 2010). The activity monitors were placed side by side under general rearing conditions (23 ± 1°C, 16:8 h L:D, 30-50% RH), and were visually separated from the rest of the room with opaque plastic to avoid disturbance of the parasitoids. The activity of each individual was recorded for 12 days, with the first (to allow time for parasitoid acclimation) and last partial days of data being removed before data analysis. The average number of beam crosses per individual per day for the 10 complete days of data was calculated. We only present average total activity levels, but we also verified that the among-day and circadian pattern of activity over the course of the parasitoids’ lives was similar among treatments (unpublished data). Patterns and causative factors related to locomotor activity variation in *T. japonicus* will be presented and discussed in detail in a separate, future study (PKA, in preparation).

To measure fecundity, we dissected *T. japonicus* from each treatment at two different time points after emergence (0 days and 12 days). Previous studies in our laboratory (see also *Results* of the current study) have shown that *T. japonicus* does not emerge with its full complement of eggs; that is, *T. japonicus* is partially synovigenic (see Jervis et al. 2001). In fact, maximum egg load in *T. japonicus* is not reached until ∼5-8 days after emergence and remains stable thereafter in the absence of hosts (unpublished data). Thus, our goal was to determine whether developing in eggs refrigerated for different durations affected both initial egg load (0 days since emergence) and maximum egg load (12 days after emergence) of *T. japonicus*. Egg load 0 days after emergence (i.e. < 24 hours since emergence) was determined by immediately freezing (−30ºC) the second female emerging from each replicate on the day of emergence and later dissecting their abdomen with fine forceps under a dissecting microscope to count the number of mature ovarioles (opaque with well-defined chorion) (see Abram et al. 2012 for more details on methodology). Egg load 12 days after emergence was determined by freezing and dissecting females used in the activity monitor experiment (see above). We have previously determined that age-specific potential fecundity in *T. japonicus* using this dissection method corresponds well to realized fecundity (i.e., when females are offered hosts) (PKA, in preparation).

To estimate longevity, the third female to emerge from each replicate was placed into a plastic vial (“Wide Drosophila vials”, diameter: 25 mm, length: 95 mm, Diamed, Canada) with drops of honey provided *ad libitum* and survival (alive or dead) was recorded three times weekly, with longevity calculated as the time difference between the midpoint between the last two survival observations and the day the parasitoid emerged.

To measure female dry body weight, a proxy for body size, the fourth through sixth emerging *T. japonicus* females from each replicate were placed in 75% ethanol in a 1.7 mL microcentrifuge tube. Next, they were placed into a glass chamber containing desiccant for 48 h. The dried parasitoids were then removed from the desiccation chamber and were weighed to the nearest µg using a ME-36S Microbalance (Sartorius, Göttingen, Germany). The three females from each replicate were weighed at the same time, and their pooled weight was divided by three to give the average dry weight for each replicate.

### Experiment 3: Comparison of refrigeration and freezing

The goal of the third experiment was to conduct a direct comparison between refrigeration and the current standard, freezing *H. halys* eggs at -80°C (McIntosh et al. 2019). More specifically, we tested whether *T. japonicus* emerging from refrigerated *H. halys* egg masses had reduced offspring survival or sublethal fitness effects compared to *T. japonicus* emerging from frozen or freshly laid *H. halys* egg masses. Freshly collected *H. halys* egg masses were glued to one quarter of a circular piece of filter paper (Sartorius 389 grade 45 mm diameter) and placed into a Petri dish (Falcon 50 mm diameter, 9 mm height). The prepared egg masses were then assigned to one of three treatments: (1) untreated (fresh eggs collected from the *H. halys* eggs from within the previous 24 h); (2) refrigerated (8°C for 14 days); (3) frozen (−80°C for 14 days). The 14-day refrigeration interval was chosen based on the results from Experiments 1 and 2 (see above) showing that this storage duration killed *H. halys* nymphs but was highly suitable for *T. japonicus* parasitism and offspring development. After the storage period, egg masses were removed from their respective treatments, allowed to reach room temperature (for refrigerated and frozen treatments), and offered individually to one female *T. japonicus* each (see above for rearing methods) for 24 h. This was conducted in three complete temporal blocks each taking place one week apart. Each temporal block consisted of 60 *H. halys* egg masses (20 from each treatment), resulting in 180 egg masses in total. Parasitoids were removed after 24 h and the parasitized egg masses were incubated under standard rearing conditions (see above). The number, development time, and sex of the emerging *T. japonicus* was recorded daily as described for Experiment 2. After parasitoid emergence, locomotor activity, fecundity (at 0 and 12 days), longevity, and weight were measured as described for Experiment 2.

### Experiment 4: Effect of post-storage conditions

The goal of the fourth experiment was to determine whether *T. japonicus* emerging from refrigerated or untreated egg masses that were then kept at different temperatures for varying time periods would have reduced offspring survival (proportion emergence) or sublethal fitness effects (development time and weight of emerging offspring). This was intended to estimate how long refrigerated egg masses, if used as sentinel egg masses for parasitoid surveys, might remain suitable after deployment, relative to viable *H. halys* eggs. We measured fewer sublethal fitness effects than in the previous experiments as activity, fecundity, and longevity are time- and labor-intensive to measure and are not directly relevant in the context of sentinel egg mass studies, which mainly aim to measure levels of parasitoid emergence.

Freshly collected *H. halys* egg masses were glued to a small piece of filter paper and placed into a Petri dish to be stored at 8°C for 14 days (the reason for the choice of duration is as described in Experiment 3). After the storage period, fresh *H. halys* egg masses were collected and, along with the refrigerated egg masses, were grouped into one of two post-storage temperature treatments: (i) “warm” temperature treatment (constant 22.9°C), or (ii) “cool” (constant 13.2°C). The warm and cool temperatures were expected to represent worst-case and best-case scenarios in the field, respectively, with the prediction that sentinel egg quality would degrade more quickly at higher temperatures (McIntosh et al. 2019). The temperature for the “warm” and “cool” temperature treatments was determined using historical data for Agassiz, British Columbia, Canada (Environment Canada 2019), calculating the mean temperature of the five days with the highest peak temperature in July (which is during the peak oviposition period of *H. halys*) and lowest minimum temperatures in September, respectively, from previous five years (2014-2018; weather.gc.ca). Post-refrigeration temperature treatments were applied by placing egg masses in 1.7 mL microcentrifuge tubes in a block heater (Henry Troemner, LLC VWR Digital 2 Block Heater 120, Model No: 949302) inside an 8°C incubation chamber and set to each of the two temperature treatments (warm or cool). Exact temperatures inside tubes were verified using a thermocouple. Egg masses were exposed to these temperatures for 1 day, 4 days, or 7 days (20 replicates per duration divided among seven temporal blocks; total n = 240). After the designated period of time at the assigned post-refrigeration temperature, each egg mass was removed from the block heater, placed into a petri dish with a drop of honey under the lid provided *ad libitum*, exposed to a female *T. japonicus* (isolated using the same method as the previous two experiments). After 24 h, each *T. japonicus* female was removed and parasitized egg masses were incubated under standard rearing conditions. Proportion emergence, development time, and sex of emerging *T. japonicus* were recorded. The weight of emerged female *T. japonicus* was measured using the methods described above.

### Experiment 5: Effects of longer-term refrigeration

The goal of the final experiment was to determine after what duration, if any, refrigeration of *H. halys* egg masses would start to negatively affect offspring survival (proportion emergence) or fitness (development time, weight) of *T. japonicus*. Again, a reduced set of sublethal effect parameters was measured in this experiment, so we did not determine the effects of longer-term refrigeration durations on parasitoid fecundity, activity, or longevity. Freshly collected *H. halys* egg masses were glued to one quarter of a circular piece of filter paper and placed into a Petri dish with water droplets under the lid to prevent desiccation. Female *T. japonicus* were isolated and prepared as described in Experiments 2, 3, and 4 above. Fresh egg masses were collected and stored at 8°C for 0 days (i.e. not refrigerated), 14 days, 21 days, 42 days, and 63 days respectively (20 egg masses per treatment divided among two temporal blocks; total n = 100). All egg masses were then exposed to an isolated female *T. japonicus* (using the same methods as in the previous three experiments) for 24 h and incubated under standard rearing conditions. Proportion emergence, development time, and sex of emerging *T. japonicus* was recorded. The weight of emerged female *T. japonicus* from each replicate was measured using the methods described above.

### Statistical analysis

For all experiments, we fit statistical models to the data that initially contained all main effects, their interactions, and used backward simplification to generate models containing only significant fixed effect factors. Analyses were done with R statistical software version 3.6.1 (R Core Team 2019). Proportion data (proportion emergence and sex ratio) were analyzed with generalized linear models (GLMs) with a binomial (if the model fit was not overdispersed) or quasibinomial (in cases of overdispersion) error distribution. Normally distributed and homoscedastic data (activity, fecundity, weight) were analyzed with linear models (LM; “lm” function). If a Bartlett’s test indicated unequal variances among treatments, a Kruskal-Wallis test was used to compare mean ranks (this only occurred once; for Experiment 3 – Weight; see *Results*). Time-to-event data (development time, longevity) were analyzed with Cox proportional hazards survival models. For experiments with paired designs, i.e. where single egg masses were split between untreated and refrigerated conditions (Experiments 1 and 2), mixed-effects versions of the above analyses were used, with the identification of each egg mass included as a random effect (“lme4” package; Bates et al. 2015). Likelihood ratio tests (“car” package; Fox and Weisberg 2019) were used to assess statistical significance (with a threshold of *p* < 0.05), except in the case of linear models or when quasibinomial error distributions were used, in which case F-tests were used for significance testing instead (Crawley 2012). Post-hoc multiple comparisons among categorical factors were done using Tukey multiple comparisons tests with the “multcomp” package (Hothorn et al. 2008). In all scatterplot figures presented in this article, the x- and y-positions of points are jittered slightly to reveal overlapping points, but models were fit to the data use the original data values. There were several missing values in many of the analyses of sublethal effects; for example, parasitoid activity, longevity, and weight could not be measured in replicates where there was no emergence or not enough emergence occurred to measure every effect. Raw data for all analyses with records of missing values are provided in the Online Supplementary Material, and final sample sizes are provided in figure captions.

## Results

### Experiment 1: Effect of short-term refrigeration on H. halys egg survival

Increasing refrigeration durations caused declines in *H. halys* egg survival (Figure 1). *Halyomorpha halys* nymph emergence was affected by an interaction between treatment type (untreated versus refrigerated eggs) and refrigeration duration (GLMM; χ^2^ = 101.64, df = 1, *p* < 0.0001). There were no changes in levels of nymph emergence in untreated eggs over the course of the experiment (χ^2^ = 1.60, df = 1, *p* = 0.21), whereas refrigeration of eggs resulted in a progressive decrease in nymph emergence over time (χ^2^ = 108.27, df = 1, *p* < 0.0001), from 82.2% nymph emergence from eggs that were not refrigerated (i.e., 0 days of refrigeration) to 0% nymph emergence after 10 days of refrigeration (Figure 1). *Halyomorpha halys* nymph emergence started to decrease markedly after 4-6 days of refrigeration (Figure 1).

**Fig. 1.**
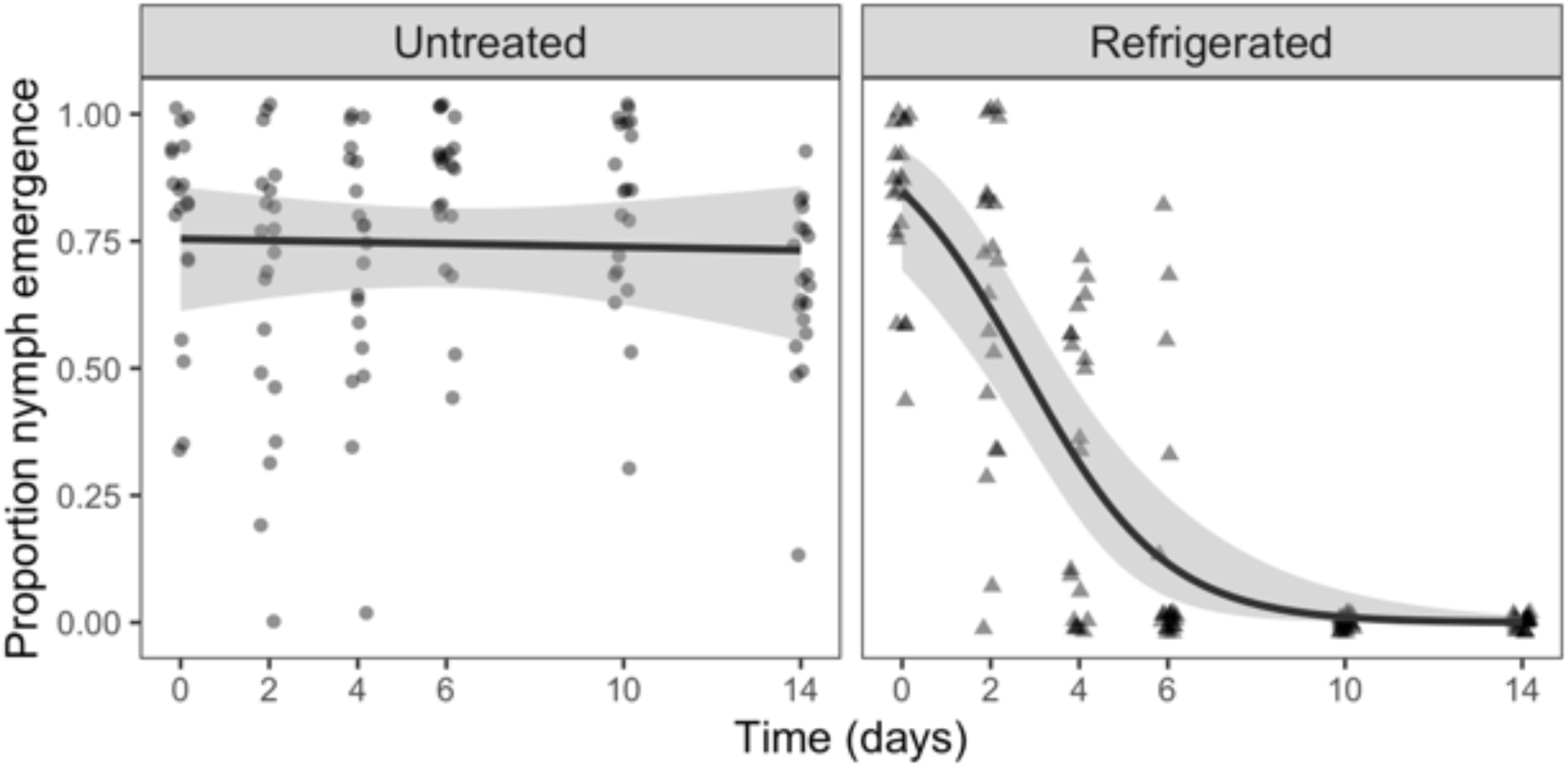
The proportion of *H. halys* nymphs emerging from untreated eggs over a 14-day period compared to paired eggs that had been refrigerated for up to 14 days (total n = 240). Trendlines are from GLMM fits considering main effects only (± 95% CIs).

### Experiment 2: Effect of short-term refrigeration of H. halys eggs on T. japonicus emergence and fitness parameters

#### Proportion Emergence

Refrigeration durations of up to 14 days did not affect *T. japonicus* emergence (Figure 2A). There was no effect of treatment type (refrigerated *vs*. untreated host eggs) (GLMM; χ^2^ = 0.69, df = 1, *p* = 0.41) or refrigeration duration (χ^2^ = 0.69, df = 1, *p* = 0.41) on the proportion of *H. halys* eggs from which *T. japonicus* emerged. The interaction between refrigeration duration and treatment type also had no effect on proportion emergence (χ^2^ = 0.019, df = 1, *p* = 0.89). There were high levels of *T. japonicus* emergence (grand mean: 95.4%) from both untreated and refrigerated eggs, even when host eggs were refrigerated for up to 14 days (Figure 2A), at which point *H. halys* eggs would all be inviable (see above).

**Fig. 2.**
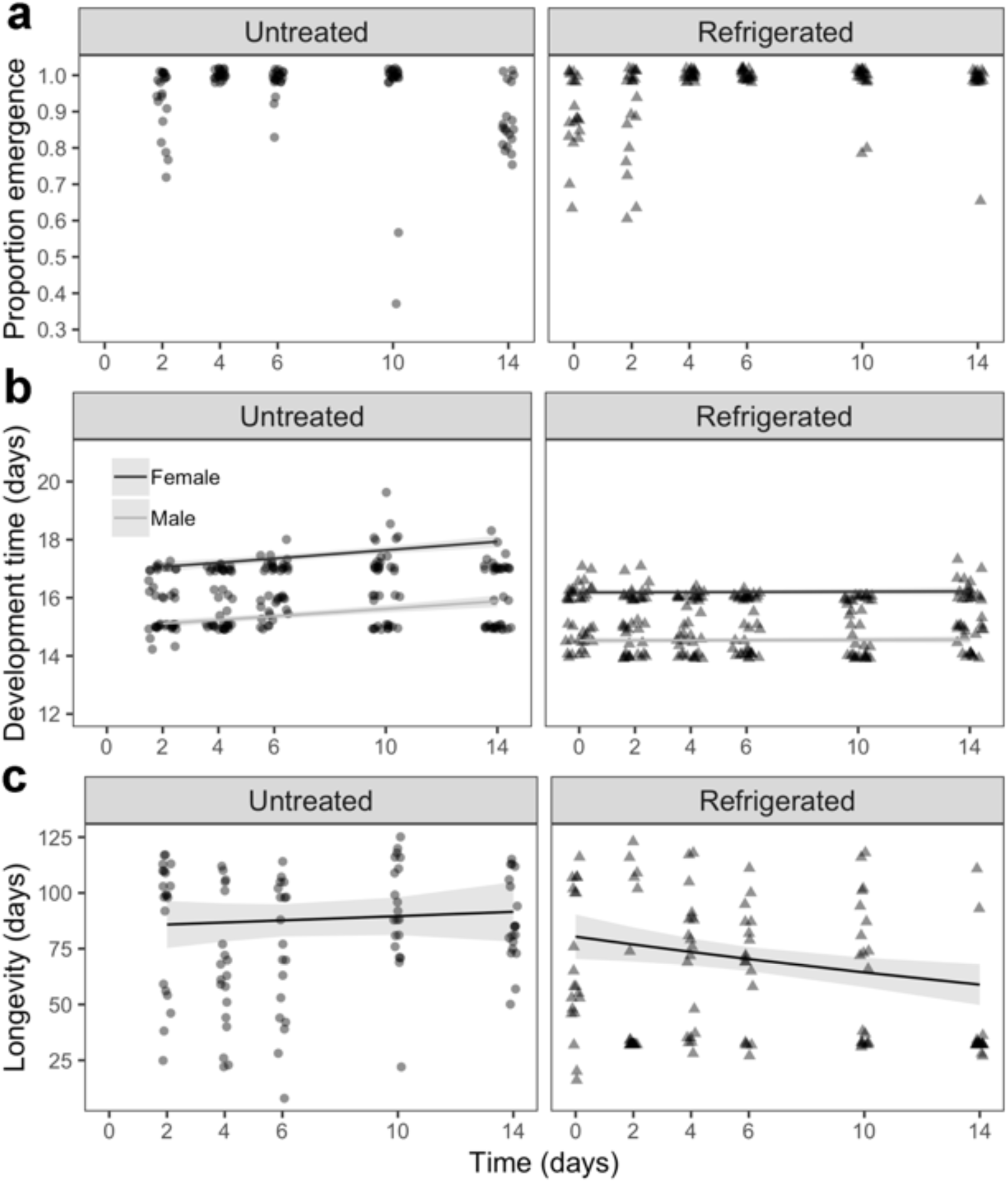
Effects of different durations of short-term *H. halys* egg refrigeration on *T. japonicus:* (a) proportion emergence (total n = 218), (b) development time, by sex (total n = 218), and (c) longevity of *T. japonicus* offspring emerging from *H. halys* eggs that were untreated, compared to paired egg masses that were collected over the same period and refrigerated for 0-14 days prior to parasitism (total n = 213). In (b) and (c), trendlines are fitted predictions from survival models (± 95% CIs).

#### Sex Ratio

Refrigeration had a small but significant effect on *T. japonicus* sex ratio (Figure S1). Refrigerated eggs, compared to fresh eggs, marginally altered *T. japonicus* sex ratio by overall increasing the percentage of male offspring from 16.4% to 21.3% on average (GLMM, χ^2^ = 4.34, df = 1, *p* = 0.037), but sex ratio was not clearly influenced by the length of refrigeration (χ^2^ = 4.34, df = 1, *p* = 0.070) or the interaction between storage length and treatment type (χ^2^ = 1.29, df = 1, *p* = 0.256).

#### Development Time

Short term refrigeration marginally decreased *T. japonicus* development time in a sex-dependent manner, but there was no clear effect of increasing refrigeration duration (Figure 2B). There was a marginal effect of the three-way interaction between treatment type (untreated *vs.* refrigerated eggs), refrigeration duration, and sex (Cox proportional hazards mixed effects model; χ^2^ = 3.83, df = 1, *p* = 0.050). Furthermore, there was an effect of the interaction between treatment type and refrigeration duration (χ^2^ = 8.97, df = 1, *p* = 0.0027) and a marginal effect of the interaction between treatment type and sex (χ^2^ = 4.08, df = 1, *p* = 0.043). For male *T. japonicus* there was an effect of treatment type on development time (χ^2^ = 107.73, df = 1, *p* < 0.0001). However, there was no effect on development time from the interaction between treatment type and refrigeration duration (χ^2^ = 0.67, df = 1, *p* = 0.41) or refrigeration duration itself (χ^2^ = 1.54, df = 1, *p* = 0.21). For female *T. japonicus* there was an effect of treatment type (χ^2^ = 148.84, df = 1, *p* < 0.0001) and the interaction between treatment type and refrigeration duration (χ^2^ = 11.40, df = 1, *p* < 0.001) on development time. However, there was no effect of refrigeration duration on development time (χ^2^ = 2.70, df = 1, *p* = 0.10). Overall, refrigeration resulted in about a one day decrease in *T. japonicus* development time compared to those developing in untreated egg masses (15.3 days *vs*. 16.4 days respectively), with *T. japonicus* females from eggs that had been refrigerated for 14 days emerging about 1.5 days earlier than those emerging from untreated eggs (16.7 days *vs*. 18.1 days, respectively) (Figure 2B).

#### Fecundity

There was no effect of refrigeration duration on the mature egg load of 0-day-old (LM; *F* = 0.02, df = 1, *p* = 0.89) or 12-day-old (*F* = 0.27, df = 1, *p* = 0.61) female *T. japonicus*. Average potential fecundity was 11.56 mature eggs for 0-day-old females, compared with 46.36 mature eggs for 12-day-old females (Figure S2).

#### Activity

There was no effect of refrigeration duration on the activity levels of female *T. japonicus* (LM; *F* = 0.38, df = 1, *p* = 0.54) (Figure S3).

#### Longevity

Increasing durations of host refrigeration prior to parasitism marginally decreased the longevity of emerging *T. japonicus* females (Figure 2C). There was an effect of the interaction between treatment type (untreated *vs.* refrigerated eggs) and refrigeration duration (Cox proportional hazards model; χ^2^ = 5.95, df = 1, *p* = 0.015). For untreated egg masses, there was no change over time in female *T. japonicus* longevity (χ^2^ = 0.71, df = 1, *p* = 0.40). However, for refrigerated egg masses there was a marginally significant negative effect of refrigeration duration in female *T. japonicus* longevity (χ^2^ = 5.16, df = 1, *p* = 0.023). Female wasps from egg masses refrigerated for 14 days (average longevity: ∼39 days) had a 42.47% decrease in longevity compared to control group female wasps (∼68 days, Figure 2C).

#### Dry Weight

There was no effect of refrigeration duration (0-14 days) on the dry weight of female *T. japonicus* (LM; *F* = 0.34, df = 1, *p* = 0.56) (Figure S4).

### Experiment 3: Comparison of refrigeration with freezing of host eggs

#### Proportion Emergence

*Trissolcus japonicus* emergence was highest from untreated and refrigerated *H. halys* eggs, and lowest from frozen eggs (Figure 3A). There was no difference in *T. japonicus* emergence between refrigerated and untreated egg masses (*p* = 0.92) (Figure 3A). There was a strong effect of treatment type (untreated/refrigerated/frozen) on parasitoid emergence (GLM; *F* = 77.74, df = 2, *p* < 0.0001), with frozen egg masses having a 58.80% and 58.24% lower proportion parasitoid emergence than untreated egg masses and refrigerated egg masses, respectively (Tukey multiple comparisons, *p* < 0.0001) (Figure 3A).

**Fig. 3.**
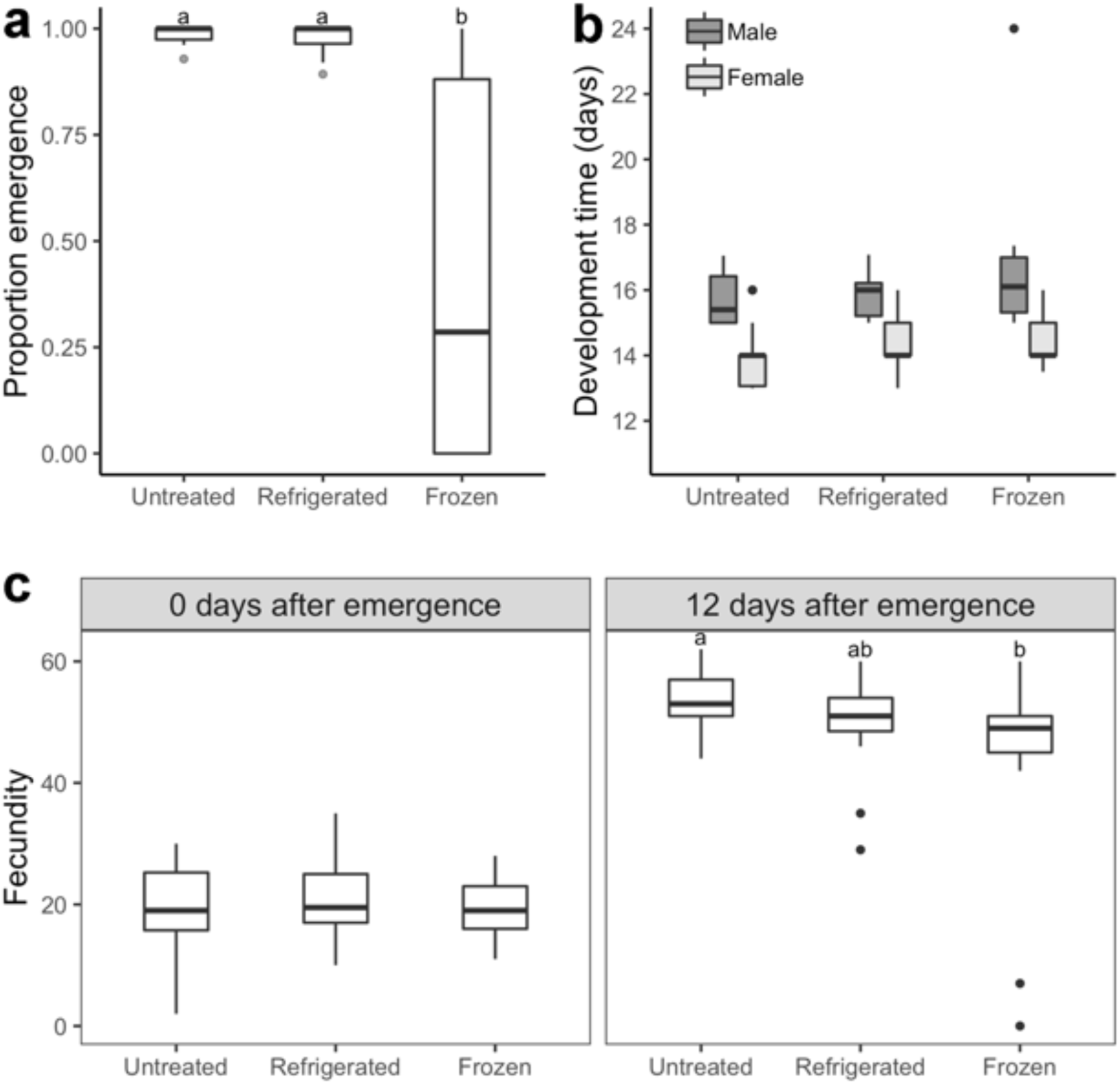
Comparison of refrigeration of *H. halys* eggs with freezing in terms of suitability for *T. japonicus*: (a) proportion emergence (total n = 90), (b) development time, by sex (total n = 120), and (c) fecundity (number of eggs determined via dissection 0 and 12 days after parasitoid emergence) of *T. japonicus* offspring developing in *H. halys* eggs that were untreated before parasitism, refrigerated (8°C) for 14 days before parasitism, or frozen (−80°C) for 14 days before parasitism (total n = 144). Within panels, boxes not labeled with the same letter are significantly different (Tukey multiple comparisons; p < 0.05)

#### Sex Ratio

There was no effect of treatment type on *T. japonicus* sex ratio (GLM; *F* = 0.21, df = 2, *p* = 0.81), which averaged 35.3% male offspring across treatments (Figure S5).

#### Development Time

Developing in previously frozen host eggs, relative to refrigerated and untreated eggs, marginally increased *T. japonicus* development time (Figure 3B). Treatment type had a marginal effect on *T. japonicus* development time (Cox proportional hazards model; χ^2^ = 6.88, df = 2, *p* = 0.032). Overall, wasps emerging from frozen eggs took about 10.6 hours (2.95%) longer to develop than wasps from untreated egg masses on average. Male *T. japonicus* developed faster than females overall (χ^2^ = 134.10, df = 1, *p* < 0.0001). There was no effect of the interaction between treatment type and sex on development time (χ^2^ = 1.07, df = 2, *p* = 0.58) (Figure 3B).

#### Fecundity

Fecundity of *T. japonicus* was reduced by developing in previously frozen eggs (relative to untreated eggs), but not by developing in refrigerated eggs, and this trend was only present for parasitoids 12 days after emergence (Figure 3C). There was no effect of treatment type on the egg load of *T. japonicus* 0 days after emergence (LM; *F* = 0.12, df = 2, *p* = 0.89). There was, however, an effect of treatment type on fecundity 12 days after emergence (LM; F = 5.47, df = 2, *p* = 0.0062). Female parasitoids from frozen egg masses had 17.22% fewer eggs than female parasitoids from untreated egg masses (Tukey multiple comparisons; *p* = 0.0041). Females from frozen egg masses had marginally (12.04%) fewer eggs at 12 days than female parasitoids from refrigerated egg masses (*p* = 0.088). However, there was no significant difference between the egg load of *T. japonicus* females from untreated egg masses and female parasitoids from refrigerated egg masses (*p* = 0.40). Averaging across treatments, potential fecundity was 19.94 eggs for 0-day-old females, compared with 50.40 eggs for 12-day-old females (Figure 3C).

#### Activity

There was no effect of treatment type on the mean activity of female *T. japonicus* (LM; *F* = 0.43, df = 2, *p* = 0.65) (Figure S6).

#### Longevity

There was no effect of treatment type on the longevity of female *T. japonicus* (Cox proportional hazards model; χ^2^ = 3.14, df = 2, *p* = 0.21) (Figure S7).

#### Dry Weight

There was more variability in dry weight of wasps emerging from refrigerated egg masses than wasps emerging from untreated egg masses (*K*^2^ = 16.39, df = 2, *p* = 0.00028). However, there was no effect of treatment type (untreated/refrigerated/frozen) on the mean ranked female parasitoid weight (Kruskal-Wallis rank sum test; χ^2^ = 3.13, df = 2, *p* = 0.21) (Figure S8).

### Experiment 4: Effect of post-refrigeration storage duration and temperature

#### Proportion Emergence

*Trissolcus japonicus* emergence remained high when developing in refrigerated *H. halys* eggs placed at cool and warm temperatures for up to 7 days following refrigeration, whereas *T. japonicus* emergence from viable host eggs declined when host eggs had been placed at warm temperatures for increasing amounts of time (Figure 4A). There was a marginal effect of the three-way interaction between temperature, length of time out of refrigeration, and treatment type (untreated *vs.* refrigerated eggs) (GLM; *F* = 2.57, df = 2, *p* = 0.079) on levels of *T. japonicus* emergence. In the warm temperature treatment, there was an effect of the interaction between refrigeration duration and treatment type (*F* = 5.38, df = 2, *p* = 0.0058). There was no effect of temperature exposure length on wasp emergence in refrigerated eggs (*F* = 0.53, df = 2, *p* = 0.59). However, proportion parasitoid emergence varied significantly among field temperature exposure lengths for untreated eggs (*F* = 13.97, df = 2, *p* < 0.0001). Untreated eggs exposed to warm temperatures for 7 days had significantly lower parasitoid emergence than untreated eggs exposed to cool temperatures after one day (24.13%) (Tukey multiple comparisons; *p* = 0.0044) and four days (21.25%) (*p* < 0.0033) (Figure 4A). In the cooler temperature treatment, there was an effect of prior refrigeration duration on parasitoid emergence (*F* = 3.34, df = 2, *p* = 0.039). However, there was no effect of treatment type (untreated *vs.* refrigerated) (*F* = 0.56, df = 1, *p* = 0.46) or the interaction of treatment type and the length of time eggs were exposed to the two temperature treatments (*F* = 0.35, df = 2, *p* = 0.71) on parasitoid emergence. Untreated egg masses exposed to warm temperatures for 7 days had 23.25% lower parasitoid emergence than their refrigerated counterparts (Figure 4A).

**Fig. 4.**
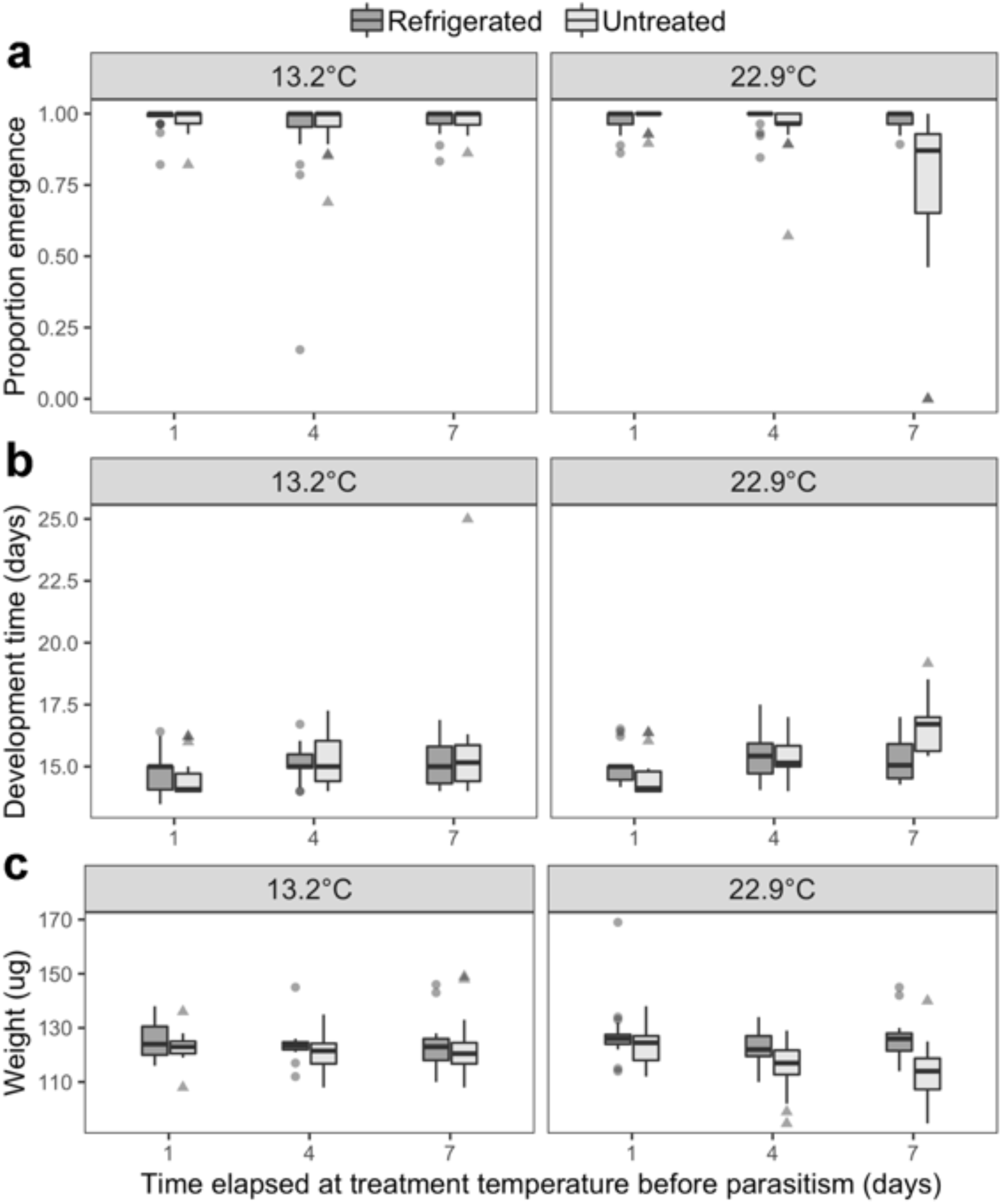
Effect of post-refrigeration temperatures on suitability of *H. halys* eggs for *T. japonicus*: (a) proportion emergence from (total n = 240), (b) development time (total n = 240), and (c) dry weight of *T. japonicus* offspring developing in *H. halys* eggs that were refrigerated (8°C) for 14 days, or untreated, and then placed at cool (13.2°C) or warm (22.9°C) temperatures for 1, 4, or 7 days before parasitism (total n = 230).

#### Sex Ratio

Sex ratio, which averaged 10.6% male, was not affected by treatment temperature (GLM, *F* = 0.12, df = 1, *p* = 0.72) or treatment type (untreated *vs.* refrigerated) (*F* = 0.057, df = 1, p = 0.81) although there was a marginal positive effect of the number of days in each treatment (*F* = 2.87, df = 2, *p* = 0.059), with the proportion of male offspring increasing slightly after 7 days for untreated eggs (Figure S9). None of the two- or three-way interactions between treatment temperature, treatment type, or treatment duration were significant (*F* < 0.68, *p* > 0.50).

#### Development Time

*Trissolcus japonicus* development time was unchanged by developing in refrigerated *H. halys* eggs placed at cool and warm temperatures for up to 7 days following refrigeration. In contrast, parasitoids developing in untreated host eggs took longer to develop when host eggs had previously been placed at warm temperatures for greater amounts of time (Figure 4B). There was an effect of the three-way interaction between treatment temperature, refrigeration duration, and treatment type on the development time of male parasitoids (χ^2^ = 6.05, df = 2, *p* = 0.048). Likewise, there was an effect of the interaction between refrigeration duration and treatment type on the development time of female parasitoids (χ^2^ = 11.37, df = 2, *p* = 0.0034). There was an effect of refrigeration duration on female parasitoid development time in untreated egg masses (Cox proportional hazards model; χ^2^ = 30.24, df = 2, *p* < 0.0001). Untreated eggs spending 7 days in either of the field temperature treatments resulted in about a 1.5 day increase in female parasitoid development time compared to spending only 1 day in either of the field temperature treatments (16.2 days *vs.* 14.8 days, respectively) (Tukey multiple comparisons; *p* < 0.001). For refrigerated eggs, there was no effect of temperature treatment, warm or cool, on female parasitoid development time (χ^2^ = 3.91, df = 1, *p* = 0.15). Refrigerated egg masses subsequently exposed to the warm temperature treatment for 7 days had a 6.44% lower development time relative to their untreated counterparts (Figure 4B).

#### Dry Weight

*Trissolcus japonicus* dry weight was unaffected by developing in refrigerated *H. halys* eggs placed at cool and warm temperatures for up to 7 days following refrigeration. In contrast, the weight of *T. japonicus* decreased as a result of developing in untreated host eggs placed at warm temperatures for increasing durations (Figure 4C). There was an effect of the interaction between treatment temperature (warm *vs.* cool) and treatment type (untreated *vs.* refrigerated) on female parasitoid dry weight (LM; *F* = 6.75, df = 1, *p* = 0.01). For warm temperature treatments, untreated egg masses produced 5.93% (7 µg) lighter females than refrigerated egg masses (*F* = 20.81, df = 1, *p* < 0.0001). Furthermore, female parasitoids from untreated egg masses exposed to the warm temperature treatment for 7 days weighed 10.62% (12 µg) less than those from previously refrigerated egg masses. Likewise, in the warm temperature treatment, there was an effect of treatment time on female parasitoid dry weight (*F* = 6.09, df = 2, *p* = 0.0031). In the cool temperature treatment, there was no effect on prior refrigeration (*F* = 1.64, df = 1, *p* = 0.20) or exposure length (*F* = 0.52, df = 2, *p* = 0.59) on female parasitoid dry weight (Figure 4C).

### Experiment 5: Effects of longer-term refrigeration

#### Proportion Emergence

With increasing refrigeration duration, up to 2 months, parasitoid emergence decreased (GLM; *F* = 22.25, df = 1, *p* < 0.0001) (Figure 5A). Refrigeration for 63 days resulted in a 9.61% decrease in the proportion of emergence compared to untreated (i.e., refrigerated for 0 days) egg masses. The largest visual decrease in proportion *T. japonicus* emergence began after 42 days of host refrigeration (Figure 5A).

**Fig. 5.**
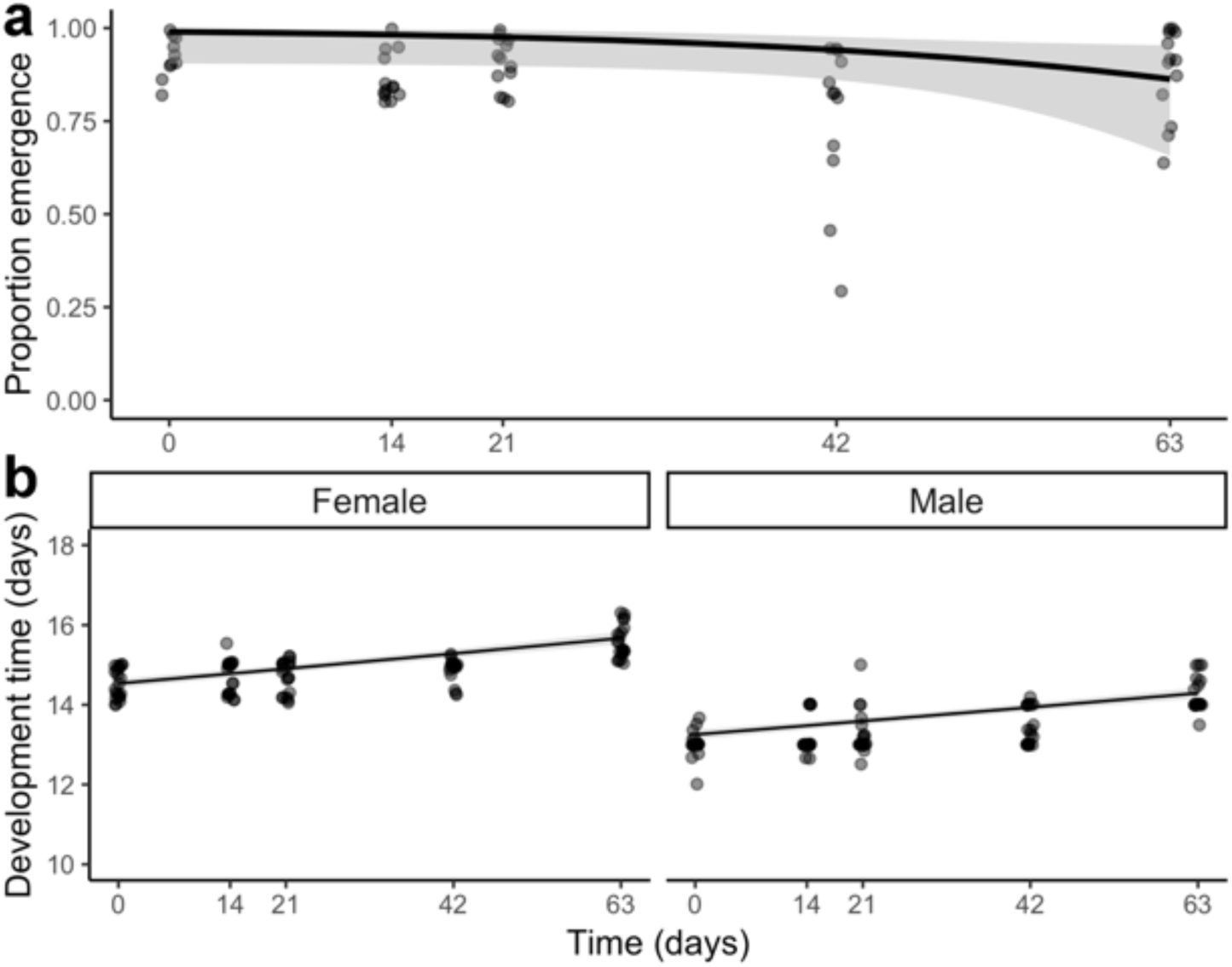
Effects of different durations of longer-term *H. halys* egg refrigeration on *T. japonicus:* (a) proportion emergence (total n = 100) and (b) development time, by sex (total n = 200), of *T. japonicus* offspring developing in *H. halys* eggs that were refrigerated (8°C) for durations of between 0 (untreated) and 63 days before parasitism. In (a) and (b), the trendlines are from a GLM with a quasibinomial error distribution and a parametric survival model, respectively (± 95% CIs).

#### Sex Ratio

There was no effect of refrigeration duration on *T. japonicus* sex ratios (*F* = 0.24, df = 1, *p* = 0.63), which averaged 12.5% male (Figure S10).

#### Development Time

Longer-term refrigeration of *H. halys* eggs prior to parasitism increased the development time of emerging *T. japonicus* (Figure 5B). There was an effect of both refrigeration duration (Cox proportional hazards model; χ^2^ = 63.58, df = 1, *p* < 0.0001) and *T. japonicus* sex (χ^2^ = 260.26, df = 1, *p* < 0.0001) on development time. However, there was no effect of the interaction between refrigeration duration and sex on development time (i.e. refrigeration duration did not affect sexes differently) (χ^2^ = 0.664, df = 1, *p* = 0.42). For emerging male parasitoids, developing in egg masses that had been refrigerated for 63 days resulted in a 1.22 day (9.35%) increase in development time. Likewise, for female parasitoids, developing in egg masses that had been refrigerated for 63 days resulted in a 1.06 day (7.30%) increase in development time. Qualitatively, most of the increase in development time appeared to occur after 42 days of host refrigeration (Figure 5B).

#### Weight

There was no effect of host refrigeration duration on female *T. japonicus* weight (LM; *F* = 0.72, df = 1, *p* = 0.40 (Figure S11).

## Discussion

Being able to store *H. halys* eggs using a method that maintains their quality as hosts would allow stockpiling of egg masses for mass rearing, monitoring and release of *T. japonicus*. The current standard for long-term storage of *H. halys* eggs is to freeze egg masses at -80ºC, which kills the host but also reduces *T. japonicus* emergence by 56-65%, with quality decreasing even more with increasing time since removal from storage (McIntosh et al. 2019). In our study, egg masses refrigerated at 8ºC for 10-14 days killed *H. halys* nymphs while also resulting in levels of parasitoid emergence (> 90%) similar to those from untreated (viable) *H. halys* eggs, with minimal (development time, sex ratio, longevity) or zero (dry weight, locomotor activity, fecundity) measurable negative sublethal effects on emerging *T. japonicus* offspring. Refrigerated egg masses also remained suitable for *T. japonicus* for longer than untreated egg masses when subsequently exposed to both warm and cool temperatures. However, some lethal and sublethal effects began to be observed when egg masses were refrigerated for more than 42 days. In the following sections, we will discuss the potential applications of the refrigeration method tested here for research on *T. japonicus* as a natural enemy of *H. halys*, relative to other options currently available for each application.

### Applications for lab rearing

*Trissolcus japonicus* laboratory colony maintenance requires a steady supply of *H. halys* eggs, the production of which is the limiting factor for maintaining and producing sufficient numbers of *T. japonicus* for adequately replicated laboratory trials and scaling up colonies for research releases. In terms of levels of parasitoid emergence and fitness parameters, refrigeration was a more effective method for short-term to medium-term stockpiling of *H. halys* eggs for *T. japonicus* mass rearing than freezing at -80ºC. Even when refrigerated for two months, parasitoid emergence was only reduced by 9.61% (compared to viable *H. halys* eggs), whereas parasitoid emergence from frozen eggs stored for 14 days was reduced by 58.80%. We expected to observe potential sublethal effects on parasitoid adults if 8ºC temperatures reduce host quality for developing immature parasitoids (Brotodjojo et al., 2006; Vinson 2010; Skillman et al. 2017; McIntosh et al. 2019). In general, sex ratio, fecundity, activity, and weight were all unaffected by short-term refrigeration of host eggs (up to 14 days) when compared to viable eggs; however, we did observe some negative effects of some refrigeration durations on longevity and sex ratio in a subset of trials. Longer-term refrigeration (up to 2 months) caused a small (1.06 day) increase in developmental time from oviposition to emergence but had no impact on sex ratio or weight of emerging parasitoids. Taken as a whole, these data support the use of refrigeration rather than freezing for short- and medium-term stockpiling *H. halys* egg masses for rearing *T. japonicus* in the laboratory. However, while we measured several life history and behavioural parameters, there may be other sublethal effects that were not measured. Additionally, informal observations show discolorations of egg masses stored for more than two months in refrigeration, which may be a visual indicator of a significant decrease in egg quality (Skillman et al. 2017). Since *H. halys* eggs stored at -80°C for more than two years still yield 25% *T. japonicus* emergence (McIntosh et al. 2019), freezing egg masses may be a better choice for long-term storage of *H. halys* egg masses for *T. japonicus* colony maintenance.

### Applications for field monitoring

The reliability of field monitoring for the presence or relative abundance of *T. japonicus* (e.g., Milnes et al. 2016; Zhang et al. 2017; Abram et al. 2019; Stahl et al. 2019a) depends on sufficient numbers and suitability of sentinel egg masses placed in the field. In earlier research on *H. halys* in invaded areas of North America and Europe, sentinel egg masses were exposed in the field after a period of freezing to allow stockpiling as well as to ensure that no additional *H. halys* were released at monitoring sites (e.g., Haye et al. 2015). Our results show that refrigeration at 8°C for 10-14 days also kills nymphs, removing this risk. Limitations of using frozen eggs include the likely underestimation of parasitism levels (Jones et al. 2014; Panizzi et al. 2018; McIntosh et al. 2019) and omission of other parasitoid-induced host mortality (Kaser et al. 2018; Stahl et al. 2019b). Consequently, some researchers have increasingly begun using viable *H. halys* eggs as an alternative (e.g., Costi et al. 2018). Refrigeration is lethal to *H. halys* eggs, and therefore, does not allow for the measurement of host mortality. However, when measuring parasitism rather than levels of host mortality is not the main goal of a survey, refrigerated eggs have the advantage of a longer period of suitability for parasitism compared to viable and frozen eggs. Our results showed that while the placement of viable *H. halys* egg masses at a relatively high average temperature (22.9ºC) for 4-7 days had a negative impact on parasitism levels and fitness parameters of emerging parasitoids, these reductions in parasitoid emergence and fitness were not observed for refrigerated eggs or in either treatment at a lower average temperature (13.4ºC). The decrease in *T. japonicus* emergence and body weight and the increase in development time with increasing physiological age of viable *H. halys* eggs is in agreement with results from other studies (Qiu 2007; Yang et al. 2018). Since the *H. halys* embryo is developing in untreated eggs (and this progresses faster at warmer temperatures), host resources and quality for immature parasitoids are likely declining (Barrett and Schmidt 1991; Vinson 2010; Skillman et al. 2017). On the other hand, eggs that have been rendered non-viable with refrigeration before nymph development progresses do not decline in quality over time (within the timespan we tested), similar to what has been found for unfertilized *H. halys* eggs (Yang et al. 2018). This is in contrast to -80ºC frozen egg masses, which are known to rapidly decline in quality for *T. japonicus* after removal from the freezer (McIntosh et al. 2019). Thus, refrigerated eggs appear to have a higher sustained quality for field monitoring of *T. japonicus* than either viable or frozen egg masses.

The relative advantages and disadvantages of using refrigerated egg masses for deployment as sentinels in field studies still need to be investigated further. High *T. japonicus* emergence levels from refrigerated eggs, when they were parasitized up to 7 days after removal from 8ºC, suggest that parasitism levels would not be underestimated if refrigerated egg masses were placed in the field and then retrieved within this period. However, whether treated eggs would be found and accepted by parasitoids at a level similar to viable sentinel eggs or naturally laid eggs in the field is unknown. One factor that could influence both host finding behaviour and host acceptance is the host cues (Conti et al. 2004; Rondoni et al. 2017; Vinson 2010; Zhong et al. 2017; Bertoldi et al. 2019) that are missing from sentinel egg masses independent of their age, viability or storage method. To include those chemical cues, one can search for naturally laid egg masses (e.g., Moonga et al. 2018); however, this is often a time-consuming and challenging process due to the cryptic nature of stink bug egg masses. Costi et al. (2018) used sleeve mesh cages with live *H. halys* adults in Italy so that egg masses laid by these adults would be associated with host cues. Unfortunately, the physical barrier may have prevented parasitoids from reaching the egg masses while simultaneously facilitating egg cannibalism by confined adults. Adding host cues by equipping *H. halys* sentinel egg masses with branches of leafy plants from stink bug rearing cages has also been attempted, but the effects on parasitism were unclear (Stahl et al. 2019c). These examples demonstrate that despite potentially missing host cues for *T. japonicus*, refrigerated *H. halys* egg masses may still be at least as effective in assessing field parasitism as current methods using viable sentinel egg masses, depending on the research question being asked.

### Applications for T. japonicus rearing to support releases

Worldwide, *T. japonicus* is being considered or has already been studied as a biological control agent; this includes its use in classical biological control as well as redistribution efforts with inoculative and inundative releases (Jentsch 2017, Charles et al. 2019; Lowenstein et al. 2020). These activities require large numbers of *T. japonicus* adults produced over short periods in order to release meaningful numbers of parasitoids throughout a growing season. For *T. japonicus* to be effective as a biological control agent in the field, maintaining parasitoid quality in terms of behavioural and physiological traits associated with host finding and parasitism success will be important (Nystrom Santacruz et al. 2017; Milnes et al. 2019; Lowenstein et al. 2019). Parameters such as locomotor activity, fecundity, and weight, may be indicative of a female egg parasitoid’s ability to locate a host, disperse, parasitize, and thus its overall efficacy as a biological control agent (Roitberg et al. 2001). Our findings support the use of refrigeration at 8°C as a stockpiling method for host eggs used to produce *T. japonicus* for releases, as most fitness-related parameters tested were not significantly different between refrigerated and untreated egg masses. In contrast, development time and fecundity were negatively affected by freezing. The notable exceptions to the general lack of negative effects of refrigeration were a ∼25% reduction in longevity for some storage durations (Experiment 2, but not Experiment 3) and a ∼1 day increase in development time (Experiment 5). The slightly increased development time in refrigerated eggs over longer durations (Experiment 5) also contradicted the finding of marginally reduced development time in refrigerated eggs over shorter durations (Experiment 2). The reason for this inconsistency is unclear. It is unknown whether these modest negative fitness effects would be enough to reduce parasitoid efficacy in the field, or whether they may be linked to other relevant life history or behavioural consequences of developing in refrigerated host eggs. It is also important to note that we did not measure some sublethal effects of refrigeration (activity, fecundity, longevity) on *T. japonicus* for refrigeration durations of more than 2 weeks. Thus, whether these longer durations would begin to have increasingly negative sublethal effects remains unclear.

With this refrigeration method, smaller rearing facilities will be capable of accumulating larger quantities of *H. halys* egg masses that can be parasitized simultaneously in support of *T. japonicus* releases. Egg masses can be stockpiled two months in advance before release; for example, to make use of egg masses produced during winter or spring *H. halys* emergence or oviposition in the field has not yet begun. Based on our measured fitness parameters, refrigeration at 8ºC appears to be an improvement over -80ºC storage for *H. halys* eggs, especially when a stable *H. halys* colony is available year-round. For facilities that wish to stockpile egg masses for maintenance or production of *T. japonicus* but do not have a stable *H. halys* colony – for example due to a high incidence of disease or fluctuating availability of technical support – freezing at -80ºC may still be a better long term-storage approach, as frozen egg masses can still be used after 2 years despite low levels of parasitoid emergence (McIntosh et al. 2019).

### Areas for future research

There are still a number of questions relating to *H. halys* egg refrigeration worthy of further study. Our study used 8ºC arbitrarily as a value between 0ºC and 12-13ºC, the threshold of freezing and the arrestment temperature of *H. halys* egg development, respectively (Nielsen et al. 2008; Haye et al. 2014), but it is still possible that other refrigeration temperatures may yield different sublethal effects. Indeed, the mechanism explaining the higher suitability of refrigerated eggs compared to frozen eggs remains unknown. We speculate that our refrigeration treatments had less impact on host quality than freezing egg masses because temperatures below 0ºC may damage host structure; for example, the chorionic envelopes or protein structures within host eggs that are important for parasitoid development (Nénon et al. 1995; Vinson 2010) by inducing ice crystal formation (Zachariassen and Hammel 1976). Refrigeration, on the other hand, may arrest host nymph development without appreciable effects, at least in the short term, on host nutritional quality.

Future studies could also address the age of the host egg when placed in refrigeration, as this might affect the method’s usefulness for stockpiling high-quality host material. In our study, we refrigerated eggs that were freshly laid (< 24 h). It is possible that egg masses placed into refrigeration that were aged for longer at room temperature would produce different results. If refrigerating 3-day old eggs yields similarly high-quality *H. halys* eggs, this would remove the need to collect egg masses at daily intervals. Further refinement to the method could also reduce the labour associated with producing host material for *T. japonicus*, and potentially extend the duration over which refrigeration retains host quality. In the future, the stockpiling of other host species or different stages of *T. japonicus* such as eggs, pupae or adults should also be considered (Orr 1988; Colinet and Boivin 2011). If refrigeration methods are broadly adopted for field use, it will be important to test whether these egg masses are attacked by native egg parasitoid species in different invaded regions and whether refrigeration makes *H. halys* eggs more or less suitable for the development of their offspring. Furthermore, if the use of refrigerated *H. halys* egg masses is successful in measuring parasitism in the field, this method could potentially be expanded to non-target native stink bug eggs for estimating levels of non-target attack by *T. japonicus* (and potentially native parasitoids as well) in the field. Studies on non-target effects on native stink bug populations that are univoltine may otherwise have difficulty in producing adequate numbers of sentinel egg masses simultaneously. If refrigerated storage effects are similar when applied to native stink bugs, this may help to expand the understanding of the impact of *T. japonicus* on such non-target species.

## Acknowledgements

We thank Jason Thiessen, Peggy Clarke, Chris Hou, Sasha Tuttle, Paul Kehler, and the Agassiz Research and Development Centre Greenhouse team for technical assistance. Thanks to Tara Gariepy, Judith Stahl, and Jake Miall for comments on an earlier version of the manuscript.

---

**Fig. S1.**
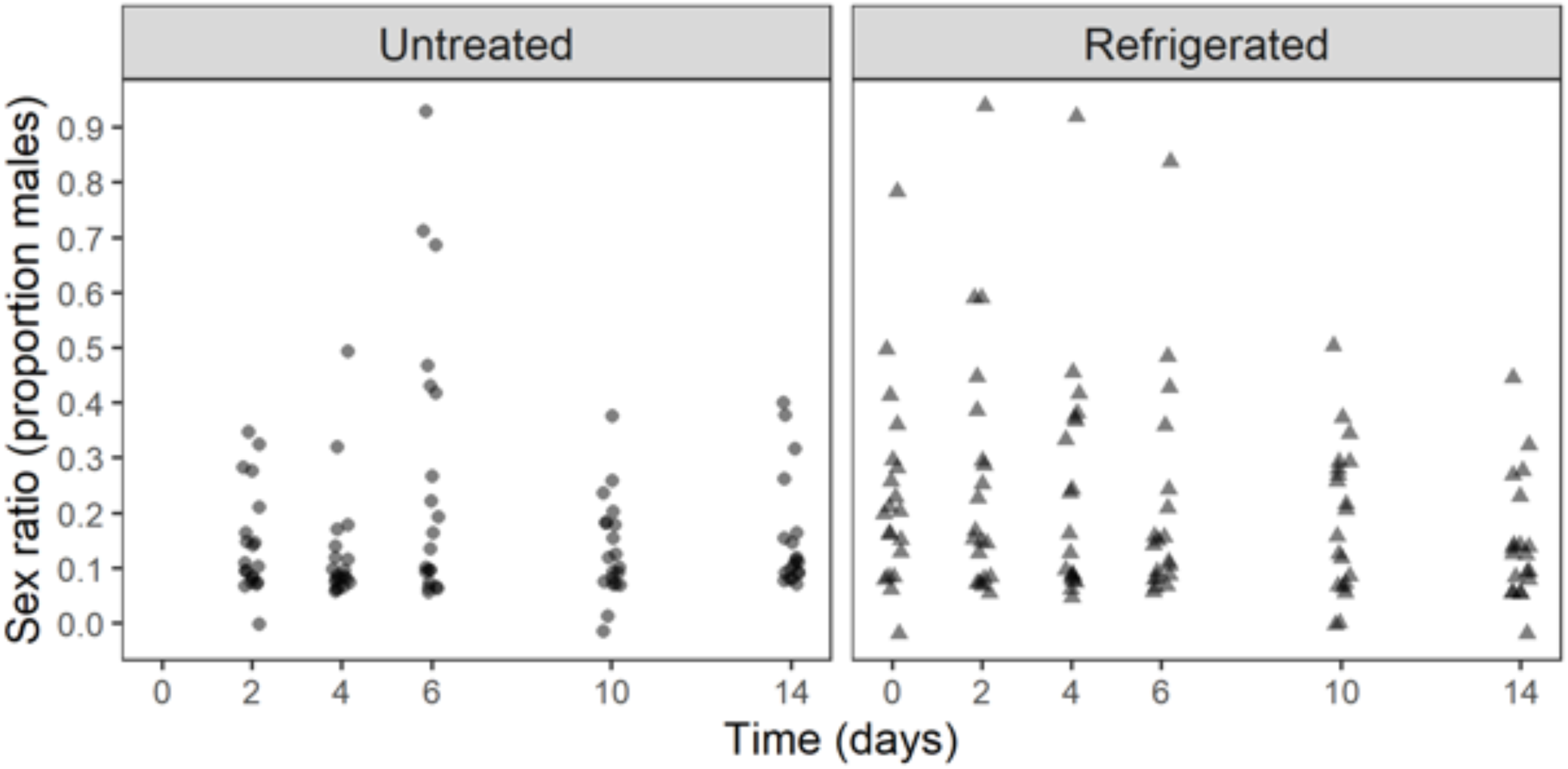
The sex ratio of *T. japonicus* offspring from *H. halys* eggs that were untreated (no refrigeration), compared to paired egg masses that were collected over the same period and refrigerated for 0-14 days prior to parasitism (Untreated: n = 99; Refrigerated: n = 98).

**Fig. S2.**
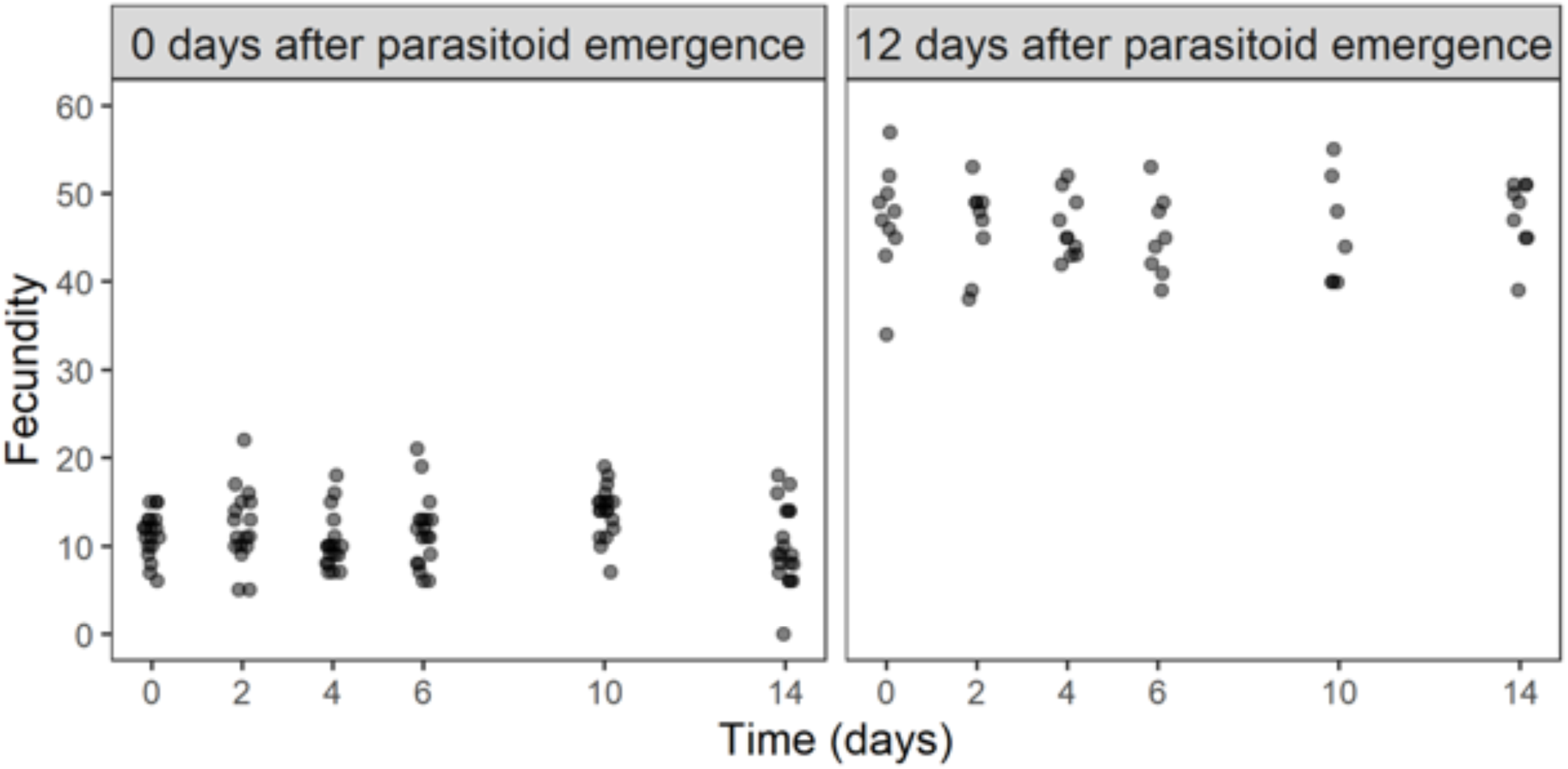
The fecundity (number of mature eggs in dissected females) of *T. japonicus* offspring emerging from *H. halys* eggs that were refrigerated for 0-14 days prior to parasitism, female parasitoid offspring that were dissected 0 days after emergence (n = 110) (left) or 12 days after emergence (n = 53) (right).

**Fig. S3.**
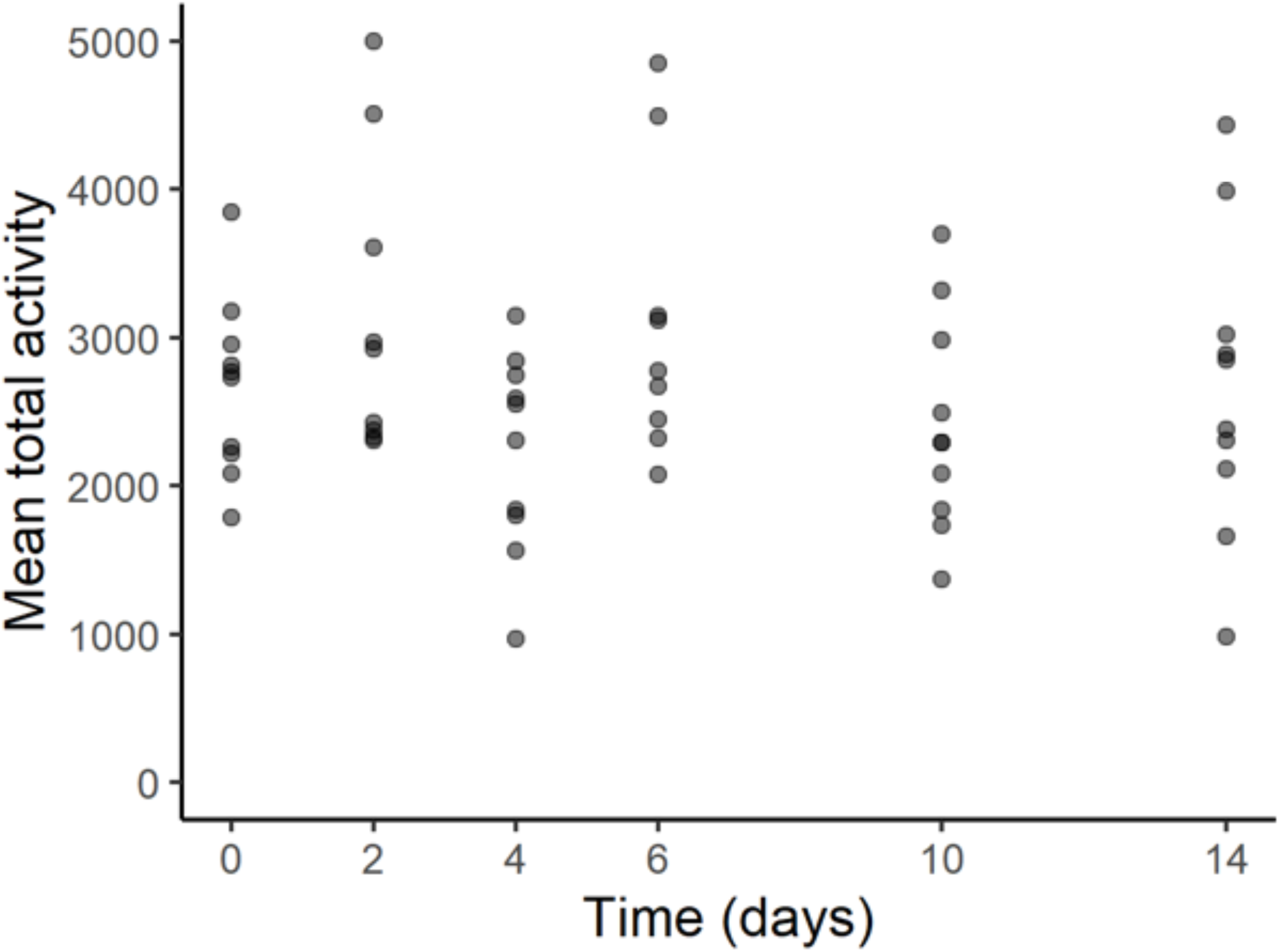
The mean total activity (average number of beam crosses per day) of *T. japonicus* adult females (averaged over the first 10 days of their lives) emerging from *H. halys* eggs that had been refrigerated for between 0-14 days prior to parasitism (total n = 58).

**Fig. S4.**
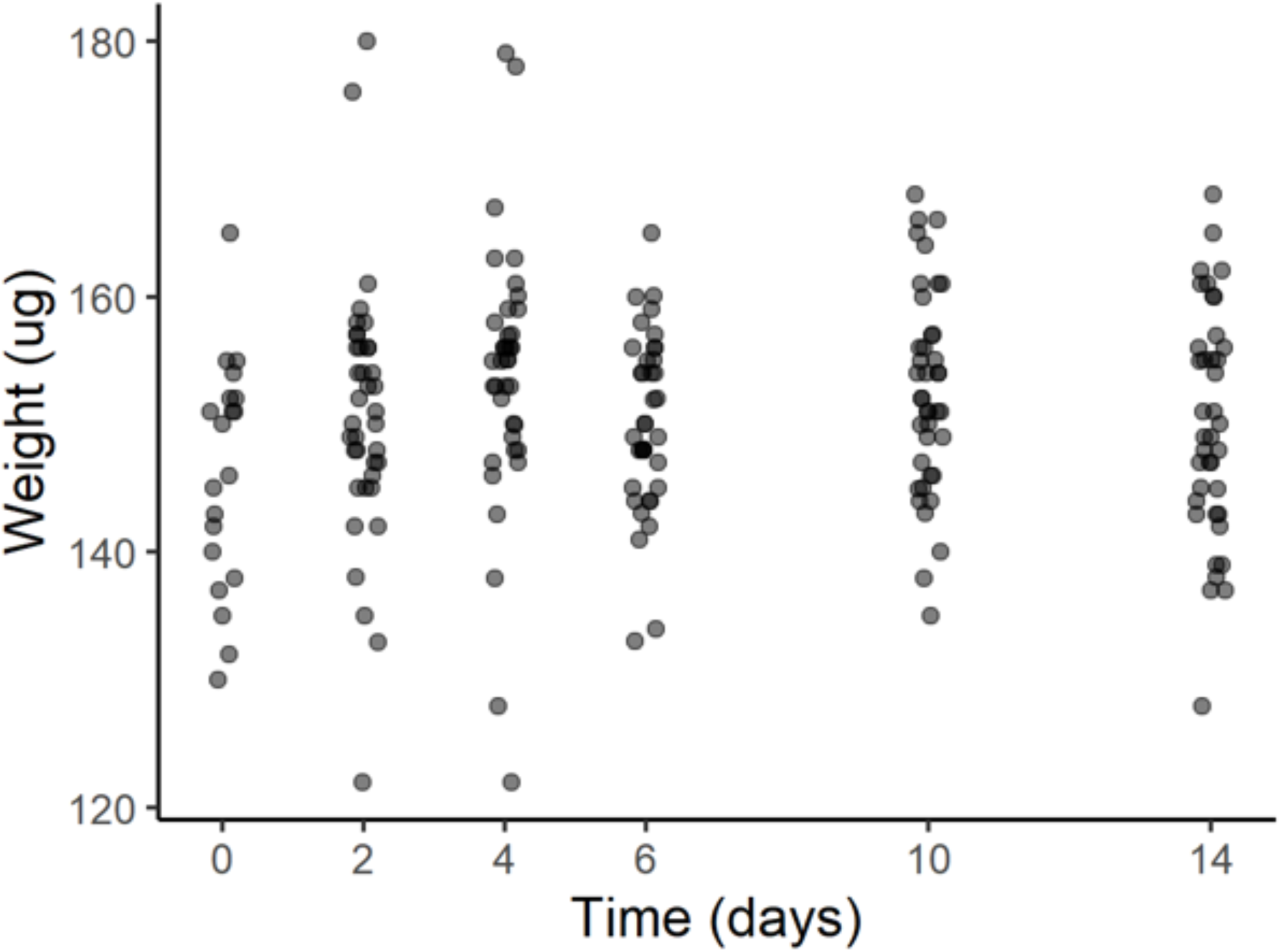
The dry weight of *T. japonicus* female offspring from *H. halys* eggs that were refrigerated for 0-14 days before parasitism (total n = 212).

**Fig. S5.**
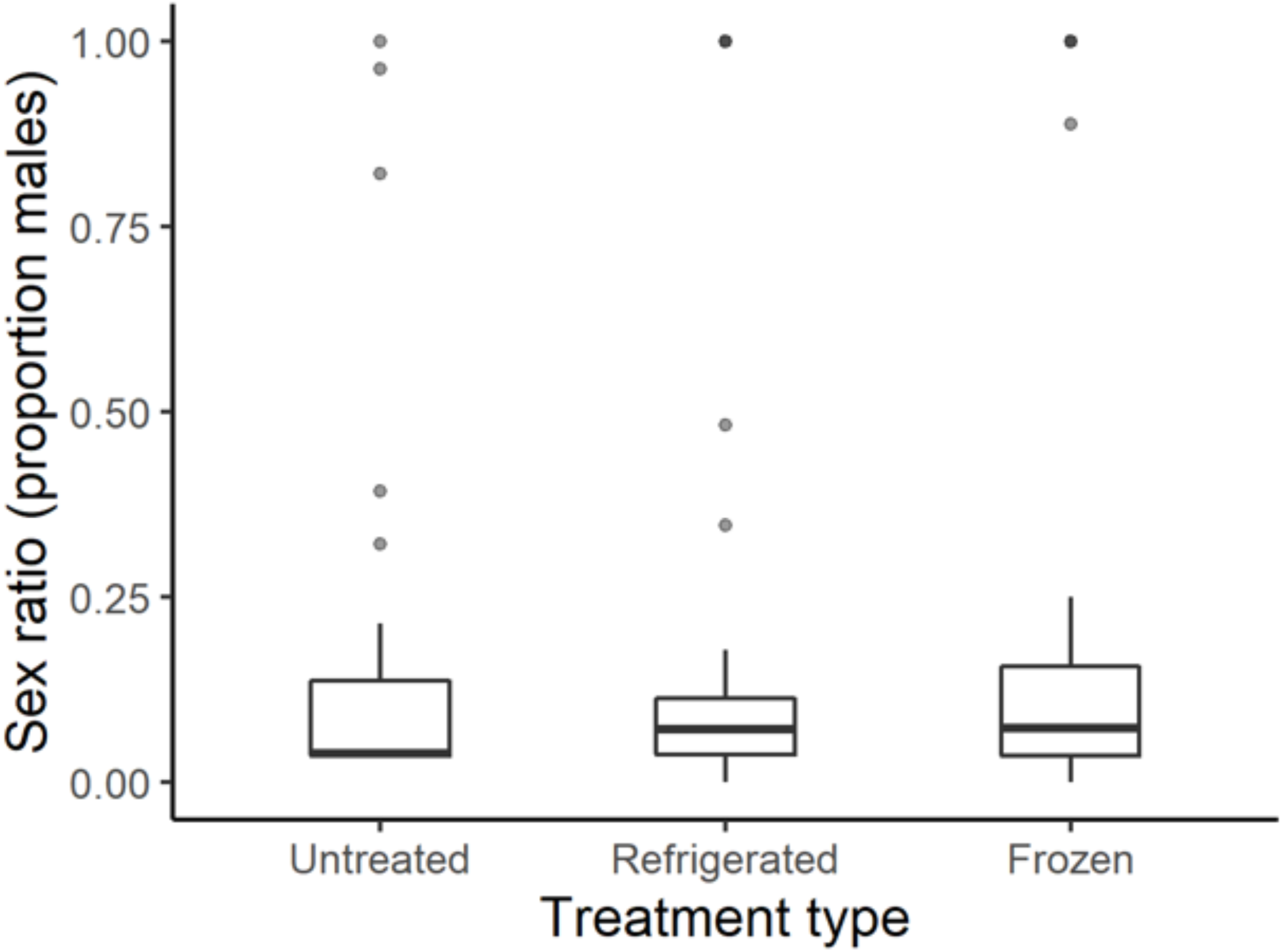
The sex ratio of *T. japonicus* emerging from *H. halys* eggs that were untreated (no refrigeration) before parasitism (n = 30), refrigerated (8°C) for 14 days before parasitism (n = 30), or frozen (−80°C) for 14 days before parasitism (n = 23).

**Fig. S6.**
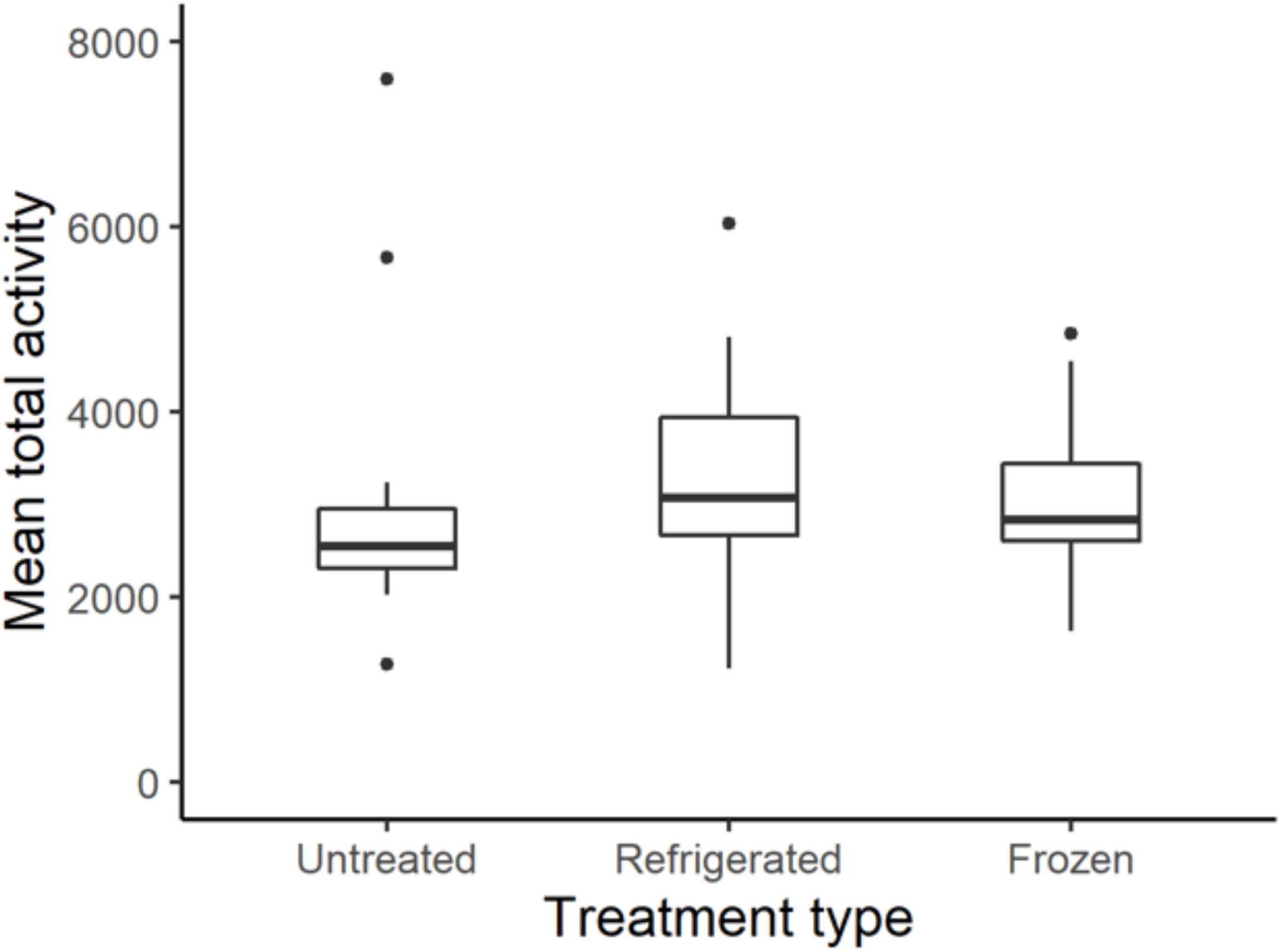
The mean total activity (average number of beam crosses per day) of *T. japonicus* adult females (averaged over 10 days of their lives) after emerging from *H. halys* eggs that were untreated (no refrigeration) before parasitism (n = 19), refrigerated (8°C) for 14 days before parasitism (n = 17), or frozen (−80°C) for 14 days before parasitism (n = 10).

**Fig. S7.**
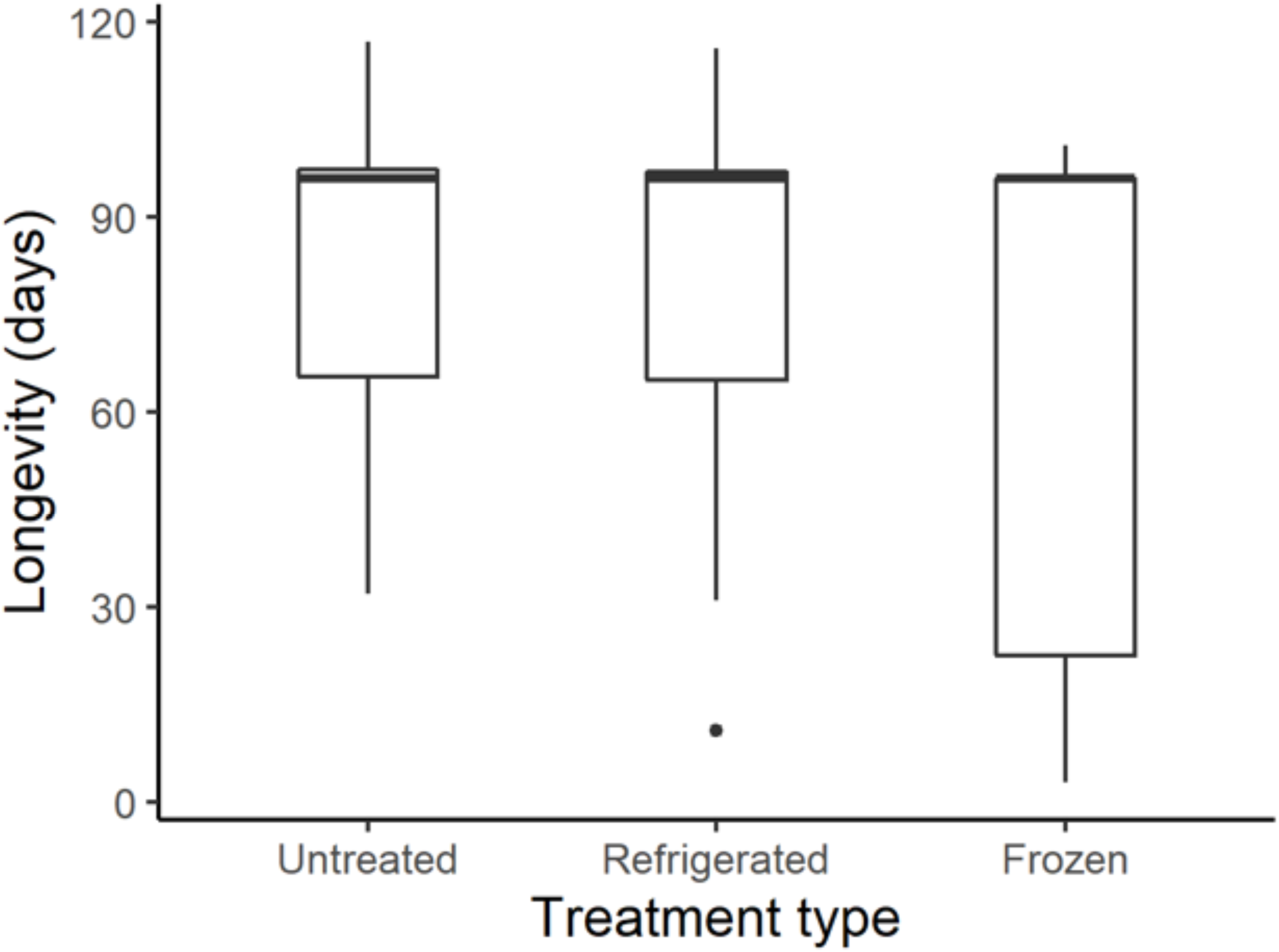
The longevity of *T. japonicus* offspring from *H. halys* eggs that were untreated (no refrigeration) before parasitism (n = 28), refrigerated (8°C) for 14 days before parasitism (n = 25), or frozen (−80°C) for 14 days before parasitism (n = 15).

**Fig. S8.**
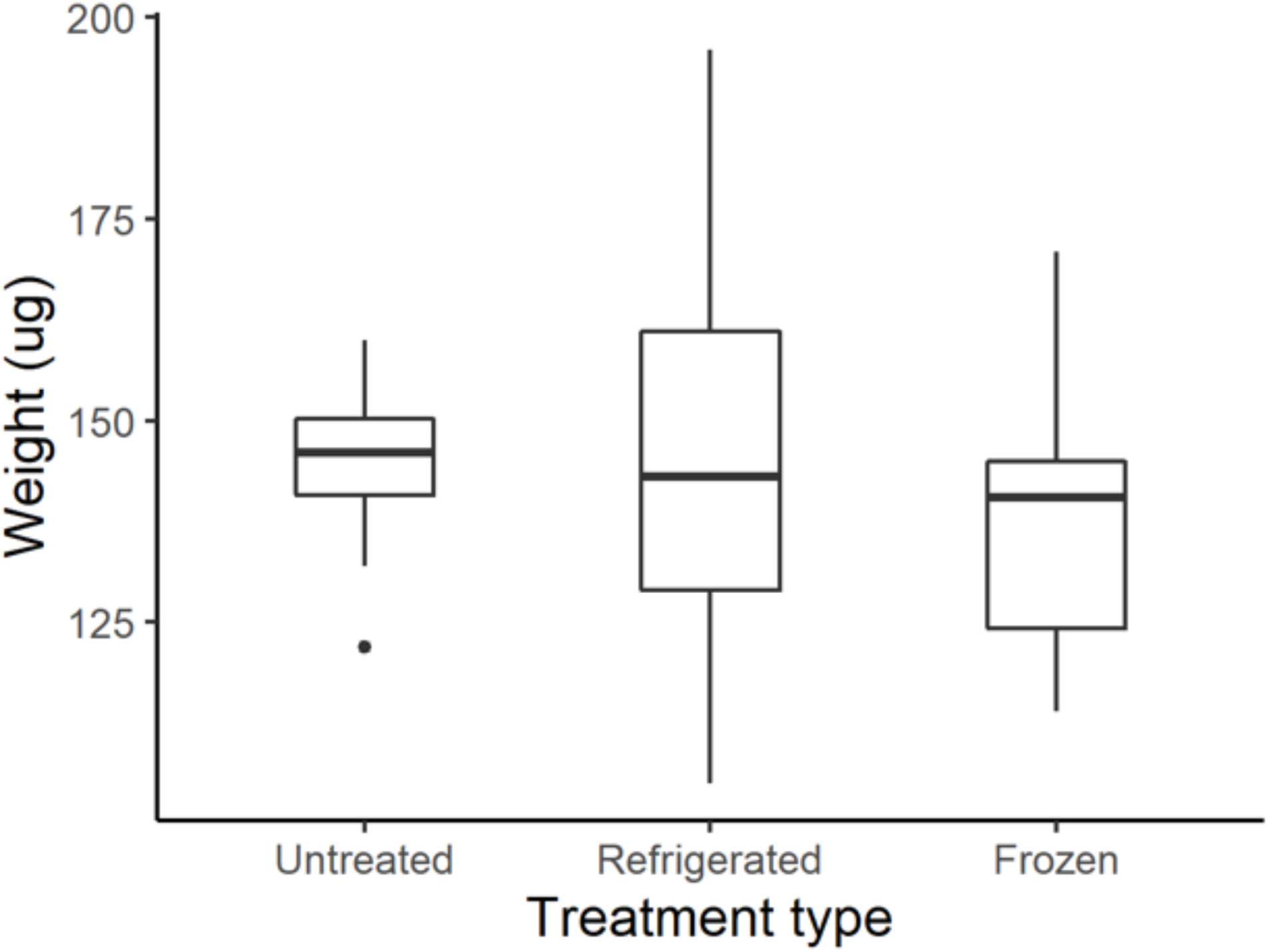
The dry weight of *T. japonicus* female offspring from *H. halys* eggs that were untreated (no refrigeration) before parasitism (n = 28), refrigerated (8°C) for 14 days before parasitism (n = 27), or frozen (−80°C) for 14 days before parasitism (n = 14).

**Fig. S9.**
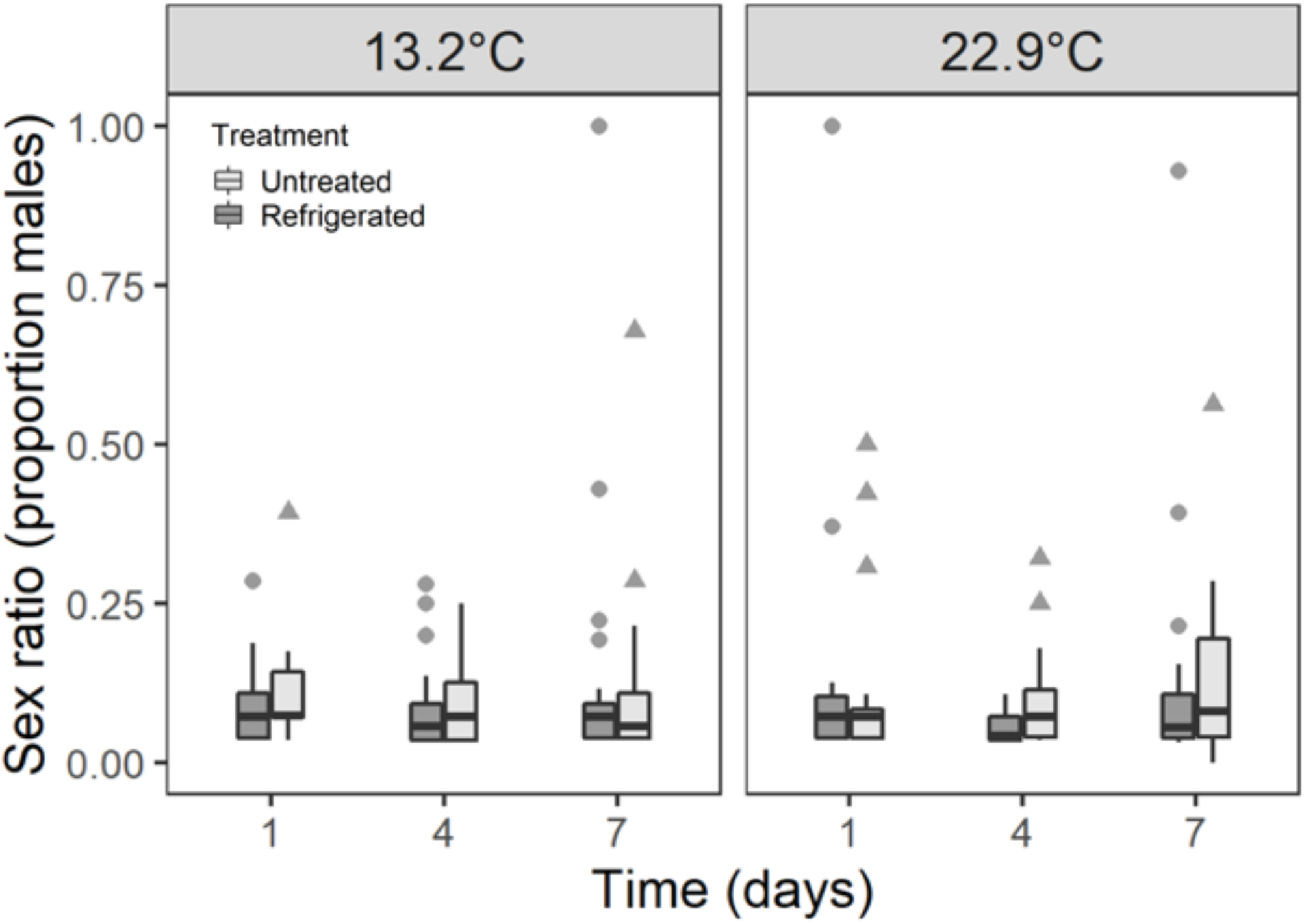
The sex ratio of *T. japonicus* offspring developing in *H. halys* eggs that were refrigerated (8°C) for 14 days, or untreated (no refrigeration), and then placed at cool (13.2°C) (untreated: n = for 1, 4, 7 day respectively; refrigerated: n = for 1, 4, 7) or warm (22.9°C) temperatures for 1, 4, or 7 days before parasitism (total n = 240).

**Fig. S10.**
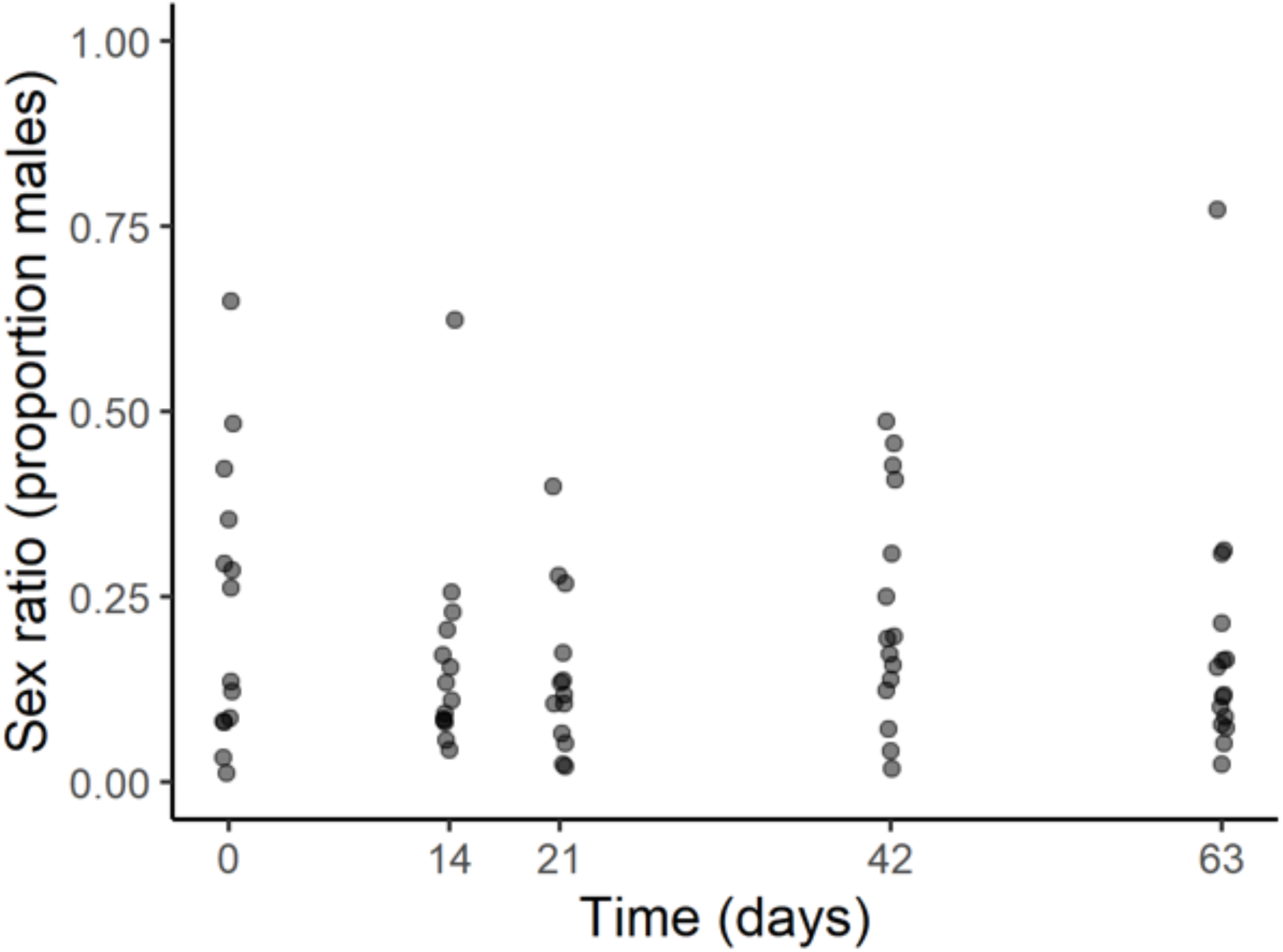
The sex ratio of *T. japonicus* female emerging from *H. halys* eggs that were cold stored (8°C) for durations of between 0 (untreated) and 63 days before parasitism (total n = 100).

**Fig. S11.**
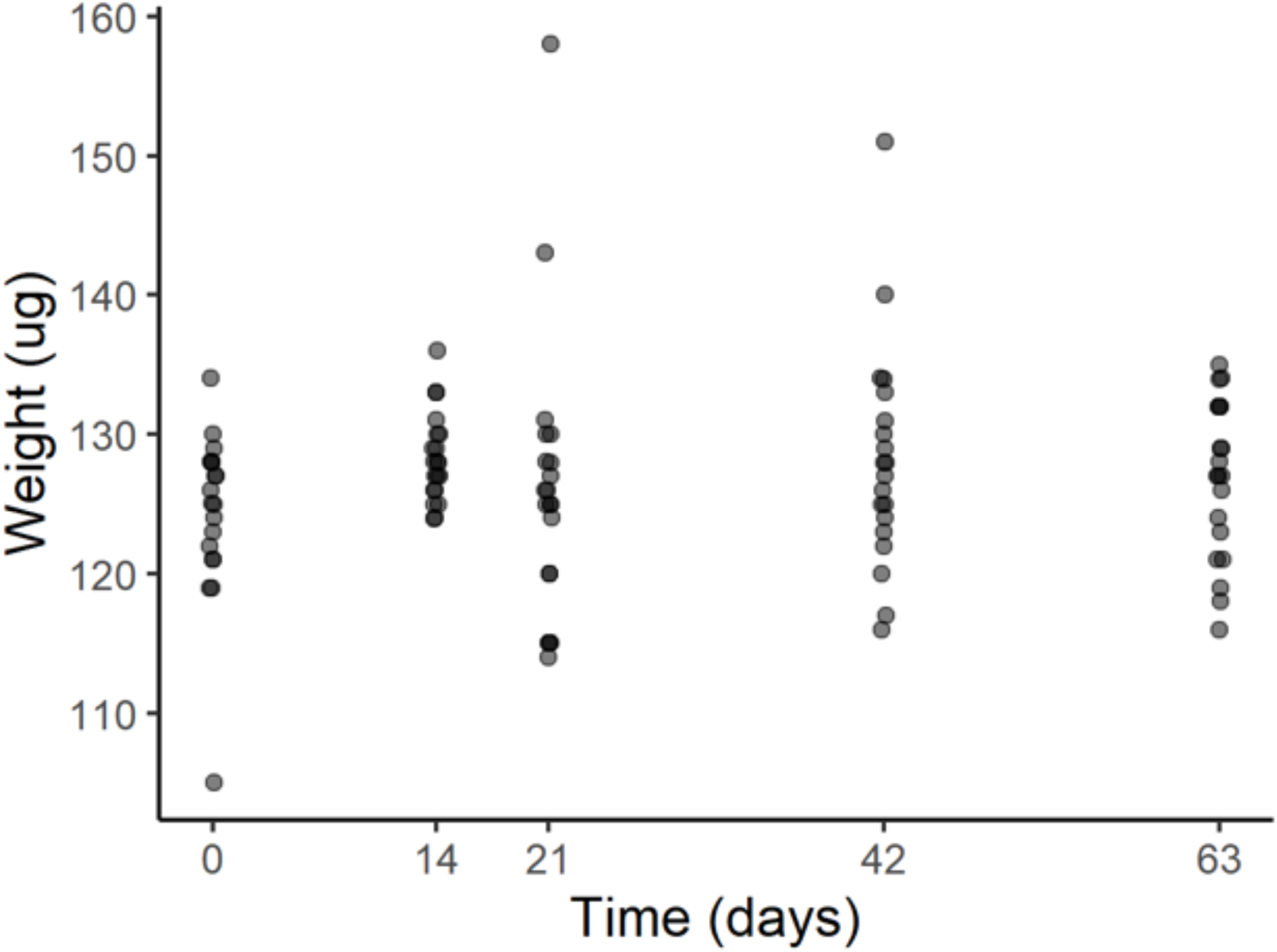
The dry weight of *T. japonicus* female emerging from *H. halys* eggs that were cold stored at 8°C for increments of between 0 (untreated) and 63 days before parasitism (n = 99).

